# Reciprocal regulation of Wnt, receptor and Dvl localizations during cell polarization

**DOI:** 10.64898/2026.11.10.765564

**Authors:** Samal Tazhibayeva, Misato Matsuo, Masayo Asakawa, Mayumi Miyazawa-Onami, Takefumi Negishi, Ariel Pani, Bob Goldstein, Hitoshi Sawa

## Abstract

Migrating cells use multiple mechanisms to interpret signaling gradients and establish polarity. However, how non-migrating cells interpret such gradients remains poorly understood. In *C. elegans*, the polarity of asymmetrically dividing seam cells is redundantly regulated by three Wnt proteins: CWN-1, CWN-2, and EGL-20. Here, we show that these Wnts, their receptors, and Dishevelled (Dvl) proteins localize asymmetrically to the posterior side of seam cells in an interdependent manner. The receptors and Dvl colocalize in Wnt-dependent, membrane-associated puncta resembling signalosomes. These findings support positive-feedback mechanisms that links Wnt localization to cell polarity. Because posteriorly expressed CWN-1 and EGL-20 have instructive roles in seam cell polarity, this mechanism may help cells interpret their gradients. In contrast, despite its anterior expression, CWN-2 localizes posteriorly, suggesting that intrinsic polarity cues also contribute to Wnt localization. Together, these findings suggest that feedback integrates Wnt gradients with intrinsic polarity cues to robustly orient cell polarity.

## Introduction

Diffusible signaling molecules can provide cells with directional information by forming spatial concentration differences across a cell or tissue. However, these differences can be shallow or noisy, whereas the resulting polarized state are often strong and persistent. How cells amplify such small differences into robust polarity is therefore a fundamental problem in cell biology. In many biological systems, feedback mechanisms are used to shape responses to signals in space and time^1^. This principle is particularly well illustrated in migrating cells, which have multiple ways to extract and reinforce directional information as they move through an external gradient. For example, shallow chemoattractant gradients are amplified intracellularly through positive-feedback interactions involving Ras, PI3K, PIP3, and F-actin in *Dictyostelium* ^2^ and PIP3, Rac, and the actin cytoskeleton in neutrophils^3^. Migrating cells can also adjust direction as they move, by comparing signaling cue over time, while continuously changing their position or shape^4–6^. In other cases, cells can modify the extracellular distribution of the signaling molecule itself, through polarized secretion or uptake, thereby generating or sharpening a gradient around themselves^7–9^. Together, these mechanisms allow migrating cells to enhance, re-sample, or even reshape the directional information available to them. Non-migrating developmental cells at a fixed position, however, are constrained in their ability to interpret a long-range diffusible signal by continuously sampling the gradient through movement. In addition, mechanisms for a stationary cell to amplify or reshape a signal gradient are unknown. How a stationary cell converts a spatial gradient into stable cortical polarity therefore remains an important question.

Wnt signaling is known to regulate the polarity of non-motile cells, primarily through the planar cell polarity (PCP) pathway, which coordinates the orientation of polarity between neighboring cells. In addition, Wnt signal gradients have been proposed to instruct the initial polarization of cells^10,11^. However, how cells interpret shallow Wnt gradients to establish polarity remains poorly understood. In both non-canonical (PCP) and canonical Wnt signaling, membrane-associated signaling complexes containing Wnt receptors and Dishevelled (Dvl) proteins play essential roles. In PCP signaling, core PCP proteins assemble into asymmetric, signalosome-like complexes at cell boundaries to establish and coordinate cell polarity^12^. In canonical Wnt signaling, Wnt promotes the formation of a receptor-associated signaling complex, or signalosome, through multivalent interactions with its receptor^13–15^, as well as polymerization of Dvl through its DIX and DEP domains^16^. However, whether these receptor–Dvl-containing complexes contribute to the spatial localization of Wnt ligands remains unknown.

In *Caenorhabditis elegans*, polarity of asymmetrically dividing somatic cells is regulated by a form of Wnt signaling called the Wnt/β-catenin asymmetry pathway^17,18^. During asymmetric divisions of six seam cells (V1-V6) that are self-renewing stem cells, Dvl proteins (DSH-2 and MIG-5) localizes to the posterior side of cells, while WRM-1/β-catenin and APR-1/APC localizes to the anterior side^19^. Although asymmetric Dvl localization is similar to PCP signaling, involvements of the core PCP components (Vangl and Prickled) and coordination of polarity among neighbors have not been reported. Seam cell polarity is regulated redundantly by three Wnt proteins: CWN-1 and EGL-20 expressed posteriorly (collectively called posterior Wnts) and CWN-2 expressed anteriorly^20^. Despite this opposing distribution, each Wnt alone can properly polarize seam cells, indicating that the spatial distribution of a Wnt ligand does not necessarily predict the direction of the resulting polarity. Consistently, reversing gradients of these Wnts can rescue polarity defects in triple Wnt mutants suggesting that polarity orientation is determined independently of Wnt gradients^21^. In contrast, in *lin-17/Fzd* mutants, reversing posterior but not anterior Wnt gradients enhances polarity reversal, suggesting that seam cells can sense posterior Wnt gradients for their polarity^17^. It is not known, however, how each cell establishes polarity in response to Wnt ligands produced at substantial distances and what is the cue for gradient-independent polarity.

To further understand the mechanism of cell polarization by Wnts, we examined the spatial organization of endogenous Wnt ligands EGL-20, CWN-1 and CWN-2, the Frizzled receptor MOM-5, the Ror receptor CAM-1, and Dvl MIG-5 on seam cell membranes. We found that all of these proteins are asymmetrically localized to the posterior side and that loss of Wnt progressively disrupts receptor polarization and reduces the membrane-associated punctate localization and colocalization of MOM-5, CAM-1, and MIG-5. Conversely, loss of CAM-1 and MOM-5 disrupts Wnt membrane localization and polarity, while loss of Dvl similarly reduces Wnt membrane localization and polarization. These reciprocal localization and polarization dependencies indicate that Wnt, receptors, and Dvl are not organized solely in a unidirectional signaling hierarchy, but mutually contribute to the spatial organization of the membrane signaling complex. Therefore, our results demonstrate that reciprocal interactions among Wnts, their receptors, and Dvl provide positive-feedback amplification mechanisms: an initial spatial bias, possibly triggered by Wnt gradients or intrinsic receptor or Dvl asymmetries, could promote nucleation of a membrane-associated complex, which would then stabilize or recruit additional components and thereby amplify the original asymmetry into robust cortical polarization.

## Results

### Wnts are localized in a gradient along seam cells

To understand how seam cells establish polarity in response to Wnts produced at a distance, we first visualized the localization of endogenous Wnts using mNeonGreen (mNG) knock-in strains: EGL-20/Wnt::mNG (*cp221*^22^), CWN-1/Wnt::mNG (*cp205;* same construct as *cp209*^23^) and mNG::CWN-2/Wnt (*os187*^17^) (Figure 1A-1C). Unlike the C-terminal fusion, the N-terminal mNG fusion of CWN-2 is functional^17^, which allowed us to follow endogenous CWN-2. Consistent with FISH analysis of Wnt transcripts^20^, mNG::CWN-2 signal was strongest in the pharyngeal isthmus, nerve ring and ventral nerve cord (Figure S1A). These tissues likely correspond to CWN-2-producing cells.

**Figure 1.**
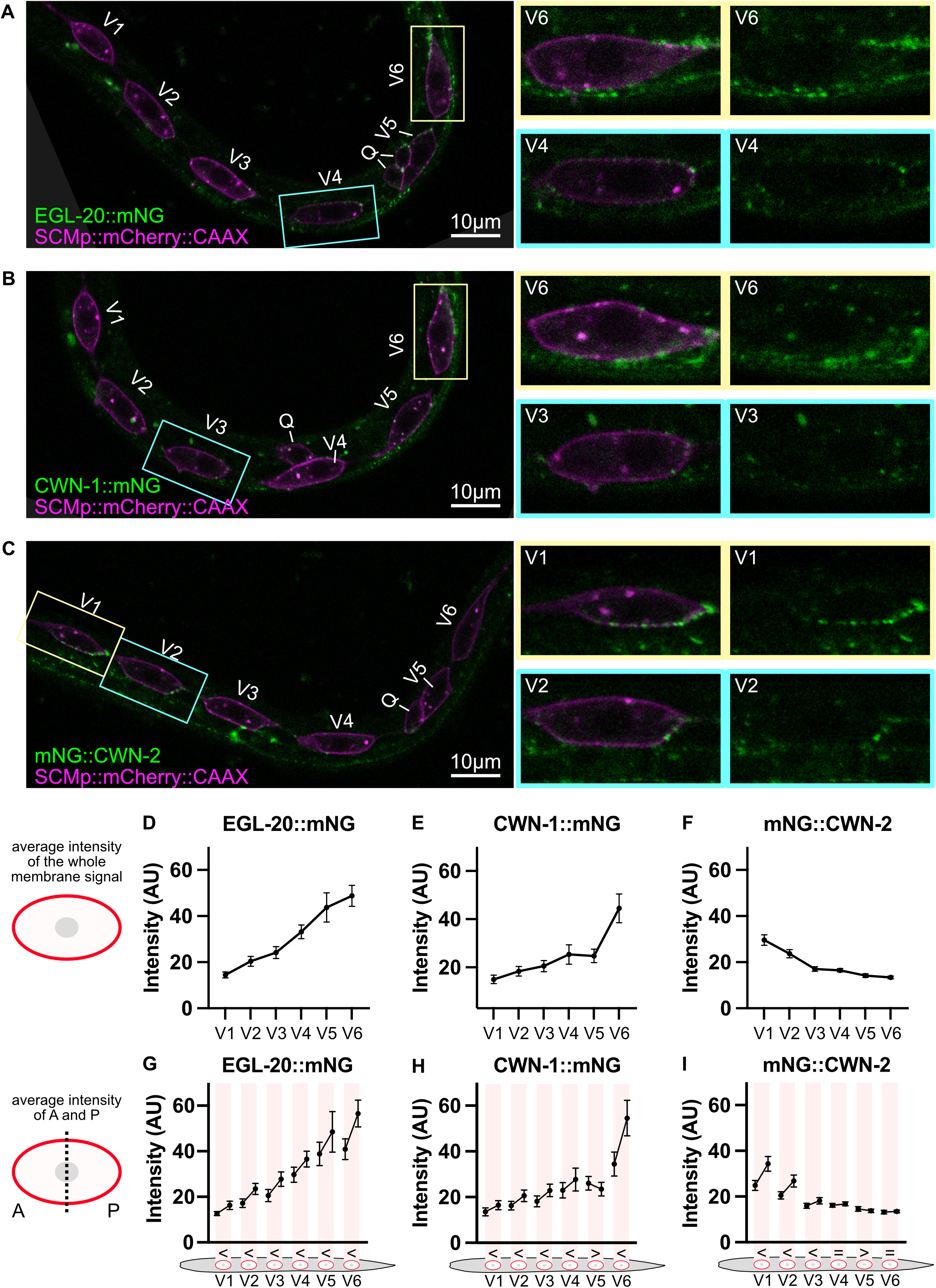
Endogenous Wnt proteins show graded and polarized localization at the seam cell membrane. **(A–C)** Representative single-plane confocal images of early L1 larvae (∼3 h post-hatching, after the initial Q cell migration and before seam cell division) expressing endogenously tagged EGL-20::mNG (A), CWN-1::mNG (B), or mNG::CWN-2 (C) (green), together with the seam cell membrane marker SCMp::mCherry::CAAX (magenta). Seam cells V1–V6 and the Q cell are labeled. Anterior is to the left. Boxed regions are shown magnified on the right. Scale bars, 10 µm. **(D–F)** Mean mNG fluorescence intensity along the whole membrane of each seam cell (V1–V6), as shown in the schematic (left), for EGL-20::mNG (D), CWN-1::mNG (E), and mNG::CWN-2 (F). **(G–I)** Mean mNG fluorescence intensity in the anterior (A) and posterior (P) halves of the membrane of each seam cell, as shown in the schematic (left), for EGL-20::mNG (G), CWN-1::mNG (H), and mNG::CWN-2 (I). Within each shaded box, the left and right points are the anterior and posterior halves of the same cell. Symbols below each cell show the result of the A–P comparison (paired t-test): “<”, A significantly lower than P; “>”, A significantly higher than P; “=”, no significant difference (P ≥ 0.05). Intensities are raw, without background subtraction. Error bars, 95% confidence intervals. Means, SD, n, and P-values for all comparisons are provided in Supplemental File 1. AU, arbitrary units.

As previously described^22,23^, all three Wnts appeared as puncta in multiple tissues, and their distribution appeared graded along the anterior–posterior (A–P) axis (Figure 1A-1C). We quantified the membrane-associated Wnt signal on V1-V6 seam cells in early L1 larvae, around 3 h after hatching, after the initial Q-cell migration and before the first seam cell division. Seam cell membranes were marked with SCMp::mCherry::CAAX (*osIs33*). Consistent with the location of their sources, mean Wnt levels on seam cells were graded. EGL-20::mNG (Figure 1D) and CWN-1::mNG (Figure 1E), which are secreted from posterior cells, decreased from posterior to anterior. mNG::CWN-2, which is secreted from anterior cells, decreased from anterior to posterior (Figure 1F).

### Wnts are localized asymmetrically on seam cells

We quantified the localization of EGL-20/Wnt::mNG, CWN-1/Wnt::mNG, and mNG::CWN-2/Wnt on anterior and posterior halves of each seam cell by determining the midpoint along the traced dorsal and ventral membranes, and mean pixel intensities were calculated for each half. Then, asymmetries of anterior/posterior signals were examined by paired t-test (numerical values for A-P comparisons of all strains are given in Supplemental File 1) as shown in Figure 1G-1I (inequality sign for significant difference and equal sign for non-significant difference). EGL-20::mNG (Figure 1G) and CWN-1::mNG (Figure 1H) showed higher signal intensity in the posterior halves of seam cells (polarized posteriorly), as previously reported for EGL-20::mNG^22^, except for V5 for CWN-1::mNG. If Wnt localization simply followed the diffusion gradients, Wnt levels should increase gradually along the seam cells. In this case, the difference within a cell (between its anterior and posterior halves) should be similar to the difference between neighboring cells (between the posterior half of one cell and the anterior half of the next). However, Wnt levels did not change continuously across cell boundaries. For EGL-20::mNG, the posterior half of one cell did not differ significantly from the anterior half of the next cell (paired t-test, p > 0.05) except for the V2–V3 boundary in which the difference ran opposite to the gradient (V2P > V3A, p = 0.024). Similarly, for CWN-1::mNG, none of the boundaries between neighboring cells among V1–V5 showed a significant difference (paired t-test, all p > 0.05). Therefore, EGL-20 and CWN-1 localization on seam cells is not a simple graded distribution.

Unexpectedly, mNG::CWN-2 was also enriched posteriorly on V1–V3 (Figure 1I). Because CWN-2 is secreted from anterior cells and graded in the opposite direction, its posterior enrichment on seam cells is counter-intuitive. Together, these results suggest that Wnt localization on seam cells is organized by seam cells, rather than simply reflecting the global Wnt gradients.

### Wnt localization and polarization on seam cells depend on their receptors

Wnt receptors are obvious candidates for mediating Wnt capture on seam cells. MOM-5/Fz, LIN-17/Fz, and CAM-1/Ror redundantly regulate seam cell polarity, as assessed by daughter cell fates^17,21^. We therefore examined how loss of these receptors affects Wnt localization on seam cells.

Mutations in Ror and Fz receptors had distinct effects on Wnt localization and polarity. In *cam-1(gm122)* mutants, all three Wnts showed decreased membrane localization in most seam cells (hereafter, “localization defects”) and defects in the asymmetry of their localization (hereafter, “polarity defects”) (Figure 2A, 2F, 2J and Table S1). In contrast, mutations in Fz receptors caused polarity defects without reducing membrane localization (Figure 2B-2C, 2G-2H and 2K-2M). In *lin-17(n3091)* mutants, both membrane localization and the asymmetry of EGL-20/Wnt::mNG were increased (Figure 2C).

**Figure 2.**
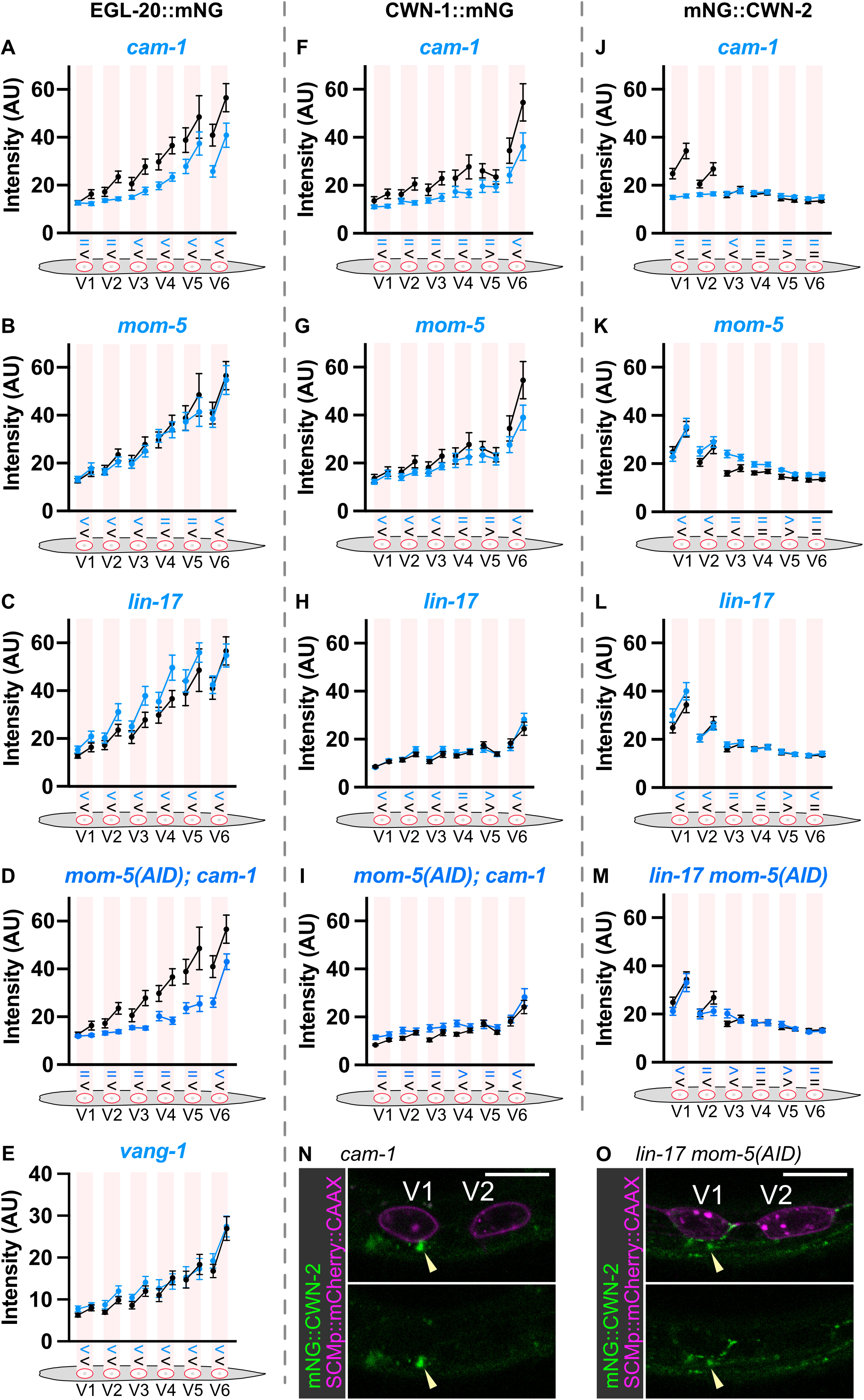
Wnt localization and polarization on seam cells depend on their receptors. **(A–M)** Mean mNG fluorescence intensity in the anterior and posterior halves of the membrane of seam cells V1–V6 for EGL-20::mNG (A–E), CWN-1::mNG (F–I), and mNG::CWN-2 (J–M) in wild-type (black) and the indicated mutant (blue) backgrounds: *cam-1* (A, F, J), *mom-5* (B, G, K), *lin-17* (C, H, L), *mom-5(AID); cam-1* (D, I), *lin-17 mom-5(AID)* (M), and *vang-1* (E). *mom-5* mutants are homozygous progeny of heterozygous mothers. Within each shaded box, the left and right points are the anterior and posterior halves of the same cell. Symbols below each cell show the result of the A–P comparison (paired t-test) for the mutant (blue, top) and wild type (black, bottom) as shown in Figure 1. Wild-type controls were imaged within 12 months of the corresponding mutant samples. **(N, O)** Representative single-plane confocal images of mNG::CWN-2 (green) and SCMp::mCherry::CAAX (magenta) in V1 and V2 of *cam-1* (N) and *lin-17 mom-5(AID)* (O) larvae. Arrowheads indicate coelomocytes. Anterior is to the left. Scale bars, 10 µm. Animals were imaged at early L1 (∼3 h post-hatching). Intensities are raw, without background subtraction. Error bars, 95% confidence intervals. Means, SD, n, and P-values are provided in Supplemental File 1. AU, arbitrary units.

The polarity defects caused by each Fz mutation affected specific seam cells. *mom-5(ne12;* homozygous from heterozygous mother) mutations disrupted the polarity of EGL-20 (Figure 2B) and CWN-1/Wnt (Figure 2G) in V4 and V5, and that of CWN-2/Wnt in V3 (Figure 2K). *lin-17* mutations affected the polarity of CWN-1 in V4 (Figure 2H) and that of CWN-2 in V3 (Figure 2L). These results might suggest that the contributions of the two Fz receptors to Wnt polarity vary among seam cells.

In *cam-1* mutants, although the overall levels of EGL-20 and CWN-1 were reduced (Figure 2A, 2F and Table S1), their mean levels showed a graded distribution across V1–V5, with significant differences between each pair of neighboring seam cells (paired t-tests, p < 0.05 for each pair; Table S2). Thus, the loss of polarized Wnt localization in *cam-1* mutants shown in Figure 2A and 2F) is unlikely to be simply a secondary consequence of reduced Wnt levels.

Remarkably, the localization defect in *cam-1* mutants was especially strong for mNG::CWN-2: its signal largely disappeared except in producing cells and coelomocytes, which take up secreted proteins from the body cavity (Figure 2J, 2N and S1B), indicating that mNG::CWN-2 is still produced and secreted. Therefore, CAM-1 is essential for CWN-2 membrane localization. However, the severe polarization defect in *lin-17 mom-5(AID)* double Fz mutants except for the V1 cell (Figure 2M and 2O), suggests that CAM-1 is not sufficient to polarize CWN-2. In this experiment, we depleted MOM-5 using the auxin-inducible degron (AID)-tagged *mom-5* allele (*os171*)^17^.

We also examined whether the core PCP component VANG-1/Vangl is involved. In *vang-1(tm1422)* mutants, the localization and polarity of EGL-20::mNG were comparable to those in wild-type animals (Figure 2E), indicating that polarized Wnt localization on seam cells does not require VANG-1. The result is consistent with nearly normal daughter cell fates in *vang-1* mutants (95% in V1 and V4 and 100% in V2, V3, V5 and V6. n=20).

Overall, Wnt polarization on seam cells depends on three receptors. Even though some of single receptor mutations disrupted Wnt polarity in specific cells, this does not necessarily mean that the other receptors do not contribute to polarity in those cells. For example, although CWN-2 polarity is significantly disrupted in *cam-1* but not *mom-5* or *lin-17* single mutants, it is also disrupted in *lin-17 mom-5(AID)* double mutants (Figure 2M). Similarly, *mom-5(AID); cam-1* strains showed more extensive polarity defects in EGL-20 (Figure 2D) and CWN-1 (Figure 2I) than each single mutant. These results indicate that the three receptors contribute redundantly to Wnt polarization. We therefore propose that three receptors have partially redundant functions in regulating Wnt polarity across seam cells, consistent with their previously reported functional redundancy in regulating daughter cell fates^17^. Even in V1, where the polarity of all three Wnts was disrupted in *cam-1* single mutants, MOM-5 may also contribute to polarity, as suggested by the more severe daughter-cell fate defects in *mom-5(AID); cam-1* double mutants (100%) than in *cam-1* single mutants (25%)^17^.

### Receptors show graded and polarized localization along seam cells

Since Wnt polarity depends on the receptors, the receptors themselves might be asymmetrically localized as reported for MOM-5/Fz in the early embryo^24^ and LIN-17/Fz in the T cell^25^ using transgenic strains that might cause overexpression. To examine the localization of endogenous receptors on seam cells, we used endogenously tagged CAM-1/Ror::mNG (*cp243*^26^) (Figure 3A) and MOM-5::mNG (*cp367*^26^) (Figure 3B), and LIN-17::mNG (*os229*; this study) (Figure 3C). Localization was quantified using a similar approach as for Wnts; however, only the dorsal side of seam cells was analyzed, because signal from the ventral side overlaps with adjacent muscle cells, making it difficult to assign receptor signal to the seam cell unambiguously. For the same reason, V4 or V5 cells contacting Q and its daughter cells were excluded from the analysis, because these cells also express the receptors.

**Figure 3.**
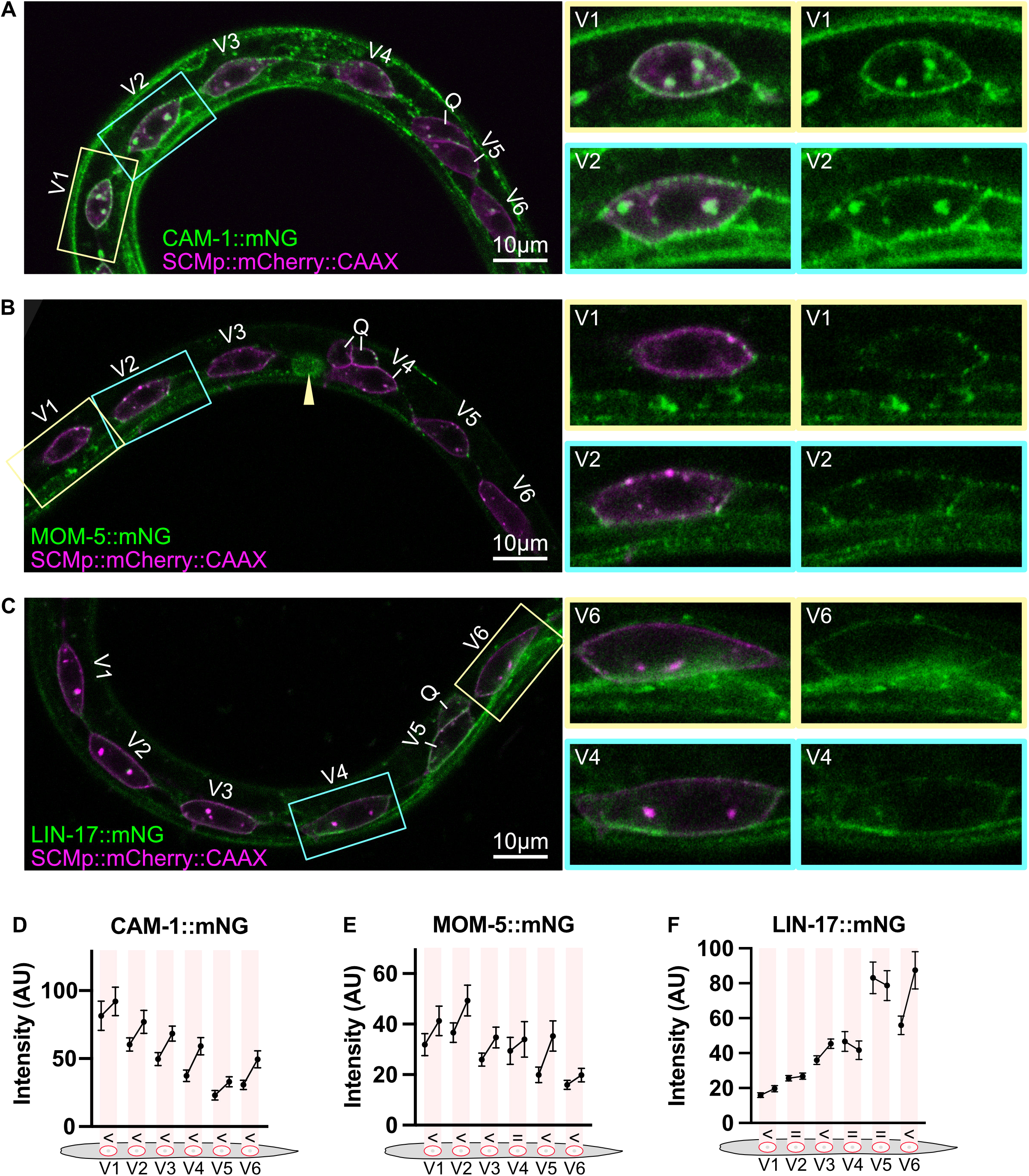
Wnt receptors show graded and polarized localization at the seam cell membrane. **(A–C)** Representative single-plane confocal images of early L1 larvae (∼3 h post-hatching) expressing endogenously tagged CAM-1::mNG (A), MOM-5::mNG (B), or LIN-17::mNG (C) (green), together with the seam cell membrane marker SCMp::mCherry::CAAX (magenta). Seam cells V1–V6 and the Q cell are labeled. Arrowhead in (B) indicates MOM-5::mNG signal in the gonad. Anterior is to the left. Boxed regions are shown magnified on the right. Scale bars, 10 µm. **(D–F)** Mean mNG fluorescence intensity in the anterior and posterior halves of the dorsal membrane of seam cells V1–V6 for CAM-1::mNG (D), MOM-5::mNG (E), and LIN-17::mNG (F). Because the receptors are also expressed in the muscle adjacent to the ventral side of the seam cells, only the dorsal membrane was quantified, and membranes overlapping the Q cell were excluded. Within each shaded box, the left and right points are the anterior and posterior halves of the same cell. Symbols below each cell show the result of the A–P comparison (paired t-test) as shown in Figure 1. Intensities are raw, without background subtraction. Error bars, 95% confidence intervals. Means, SD, n, and P-values are provided in Supplemental File 1. AU, arbitrary units.

Consistent with previous reports for CAM-1^27^ and LIN-17^21,27,28^, all three receptors showed graded localization along the A–P axis: CAM-1::mNG (Figure 3D) and MOM-5::mNG (Figure 3E) showed anterior-high gradients, whereas LIN-17::mNG showed a posterior-high gradient (Figure 3F). MOM-5::mNG and CAM-1::mNG showed punctate localization similar to Wnts and were strongly polarized to the posterior in all seam cells (Figure 3A-3B and 3D-3E). Although LIN-17::mNG was posteriorly polarized in some cells, it did not show punctate localization unlike Wnt localization (Figure 3C and 3F). Additionally, since disruption of both *mom-5* and *cam-1* was sufficient to impair Wnt polarity (Figure 2D and 2I), subsequent analyses are focused on localization of these two receptors.

### Receptor localization and polarization depend on Wnts

In the T cell, that is the most posterior seam cell in the tail, LIN-44/Wnt determines the polarized localization of LIN-17/Fz^25^. However, LIN-44 is produced by cells adjacent to the T cell and appears to act as a short-range signal, since *lin-44* mutation does not affect daughter fates of V1-V5 in otherwise wild-type or compound Wnt mutant backgrounds^21^. Therefore, we examined whether Wnts produced at a distance can also regulate receptor asymmetry on seam cells by analyzing receptor localization in Wnt mutant backgrounds. The results show that receptor asymmetry is redundantly regulated by three Wnt genes. Single Wnt mutants (*egl-20*/Wnt*(n585)*, *cwn-1/*Wnt*(ok546)*, *cwn-2/*Wnt*(ok895)*) disrupted the asymmetry of CAM-1/Ror::mNG (Figure 4A-4C) and MOM-5/Fz::mNG (Figure 4G-4H) in some posterior seam cells. Double mutants lacking both posterior Wnts, *cwn-1; egl-20*, showed more pronounced asymmetry defects in both receptors, affecting seam cells in the posterior and middle parts of animals (V3–V6), with reversed polarity in V5, consistent with strong V5 polarity reversal *cwn-1; egl-20* assessed by cell fates^21^. In anterior seam cells (V1–V2), receptor localization remained asymmetric (Figure 4D and 4I). In triple Wnt mutants, however, asymmetric localization of MOM-5::mNG was mostly abolished (Figure 4J) (residual asymmetry in V3–V4 is likely due to gonadal signal occasionally visible between V3 and V4 (Figure 3B)), indicating that CWN-2 alone, despite its opposite gradient, is able to polarize MOM-5 to the posterior in the absence of CWN-1 and EGL-20. In contrast, CAM-1::mNG retained asymmetric localization in V2–V3 even in the absence of all three Wnts (Figure 4E-4F).

**Figure 4.**
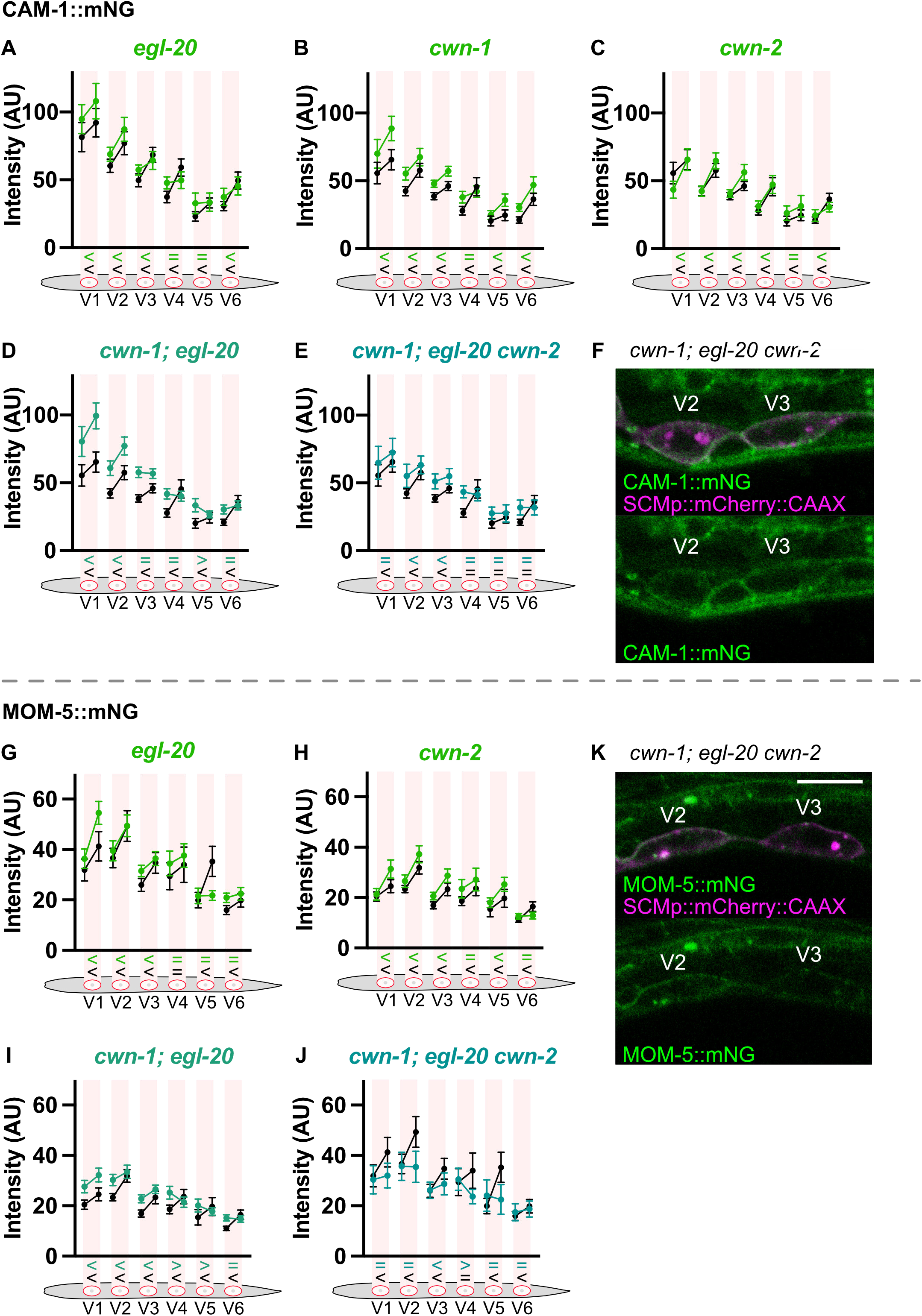
Receptor localization and polarization on seam cells depend on Wnts. **(A–E)** Mean CAM-1::mNG fluorescence intensity in the anterior and posterior halves of the membrane of seam cells V1–V6 in wild-type (black) and the indicated mutant (colored) backgrounds: *egl-20* (A), *cwn-1* (B), *cwn-2* (C), *cwn-1; egl-20* (D), and *cwn-1; egl-20 cwn-2* (E). **(F)** Representative single-plane confocal images of CAM-1::mNG (green) and SCMp::mCherry::CAAX (magenta) in V2–V3 in *cwn-1; egl-20 cwn-2* triple Wnt mutants. **(G–J)** Mean MOM-5::mNG fluorescence intensity in the anterior and posterior halves of the membrane of seam cells V1–V6 in wild-type (black) and the indicated mutant (colored) backgrounds: *egl-20* (G), *cwn-2* (H), *cwn-1; egl-20* (I), and *cwn-1; egl-20 cwn-2* (J). **(K)** Representative single-plane confocal images of MOM-5::mNG (green) and SCMp::mCherry::CAAX (magenta) in V2–V3 in *cwn-1; egl-20 cwn-2* triple Wnt mutants. In (A–E) and (G–J), only the dorsal membrane was quantified and membranes overlapping the Q cell were excluded. Within each shaded box, the left and right points are the anterior and posterior halves of the same cell. Symbols below each cell show the result of the A–P comparison (paired t-test) for the mutant (colored, top) and wild type (black, bottom) as shown in Figure 1. Wild-type controls were imaged within 12 months of the corresponding mutant samples. In (F) and (K), anterior is to the left. Scale bars, 10 µm. Animals were imaged at early L1 (∼3 h post-hatching). Intensities are raw, without background subtraction. Error bars, 95% confidence intervals. Means, SD, n, and P-values are provided in Supplemental File 1. AU, arbitrary units.

In addition, in posterior Wnt double mutants, where CWN-2 is the only remaining Wnt, receptor signal intensity was significantly increased in the anterior half of the animals (Figure 4D, 4I, Table S3), whereas in triple Wnt mutants, it was comparable to the control in CAM-1::mNG and decreased in MOM-5::mNG (Figure 4E-4F, 4J-4K and Table S3).

Overall, these results show that receptor polarity is regulated by Wnts. Taken together with the dependence of Wnt localization on receptors (Figure 2), these results suggest reciprocal regulation between Wnts and their receptors: Wnts contribute to the polarization of receptor localization, while receptors, in turn, capture and polarize Wnts on seam cells. Such bidirectional interactions could generate a positive feedback loop that amplifies and maintains asymmetric membrane localization.

### Asymmetric localization of CWN-2 can be established independently of posterior Wnts

One possible reason for the posterior polarization of CWN-2/Wnt is that it is captured by receptors polarized by posterior Wnts, even though this possibility is unlikely, because CWN-2 alone, in *cwn-1/*Wnt*; egl-20/*Wnt double mutants, is able to polarize MOM-5/Fz to the posterior (Figure 4I and 4J). To directly exclude this possibility, we examined mNG::CWN-2 localization in posterior Wnt double mutants. In *cwn-1; egl-20(os214)* double mutants, mNG::CWN-2 remained enriched at the posterior side of seam cells (Figure 5A). Therefore, the posterior polarization of CWN-2 does not require posterior Wnts. Notably, the overall mNG::CWN-2 signal on the membrane was increased in this mutant background, similar to that of receptors (Figure 4D, 4I and Table S3). One possible explanation is that, in the absence of posterior Wnts, CWN-2 has more chance to bind to receptors, which in turn may stabilize receptors on the membrane.

**Figure 5.**
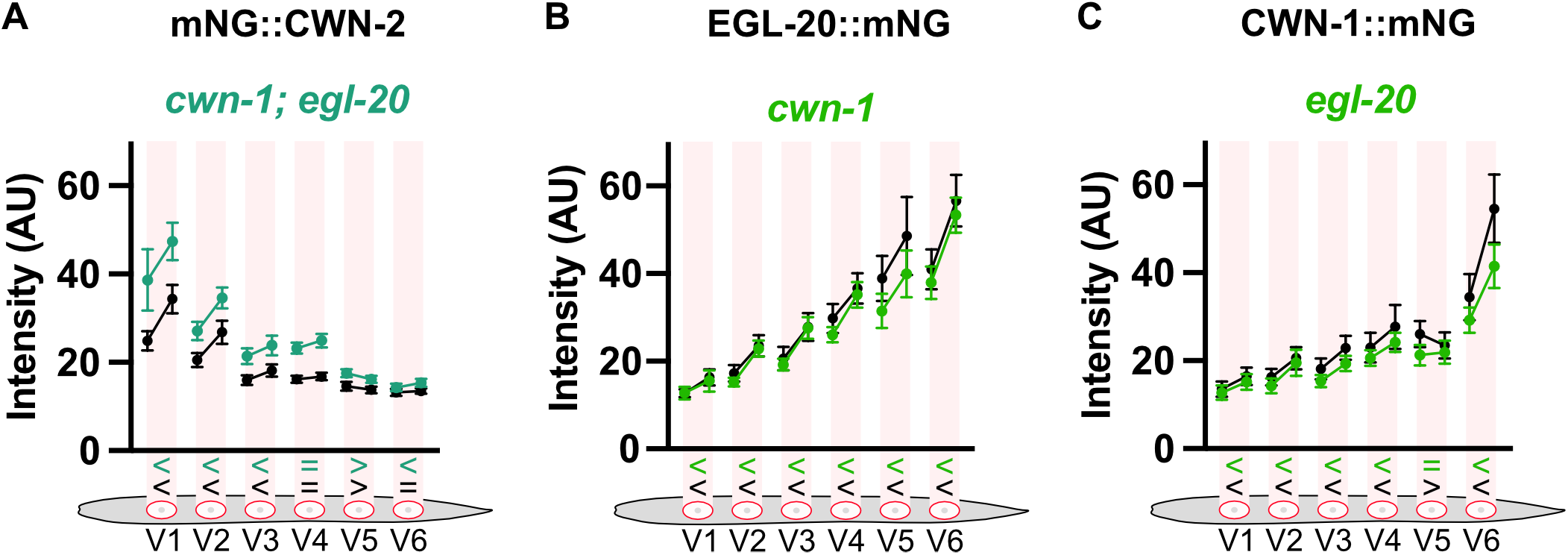
Asymmetric localization of CWN-2 can be established independently of posterior Wnts. **(A–C)** Mean fluorescence intensity in the anterior and posterior halves of the membrane of seam cells V1–V6 for mNG::CWN-2 in *cwn-1; egl-20* double mutants (A), EGL-20::mNG in *cwn-1* mutants (B), and CWN-1::mNG in *egl-20* mutants (C) (colored), compared with wild type (black). Within each shaded box, the left and right points are the anterior and posterior halves of the same cell. Symbols below each cell show the result of the A–P comparison (paired t-test) for the mutant (colored, top) and wild type (black, bottom) as shown in Figure 1. Wild-type controls were imaged within 12 months of the corresponding mutant samples. Animals were imaged at early L1 (∼3 h post-hatching). Intensities are raw, without background subtraction. Error bars, 95% confidence intervals. Means, SD, n, and P-values are provided in Supplemental File 1. AU, arbitrary units.

We also tested whether posterior Wnts contribute to each other’s asymmetric localization. EGL-20::mNG localization and asymmetry were not affected in *cwn-1* mutants (Figure 5B). Similarly, CWN-1::mNG localization and asymmetry in *egl-20(n585)* mutants were comparable to those in wild-type animals (Figure 5C), except for V5. The absence of Wnt localization defects in single Wnt mutants, despite the mild reduction in receptor asymmetry (Figure 4), is likely due to the redundancy of receptor functions.

Overall, both the anterior Wnt CWN-2 and the posterior Wnts EGL-20 and CWN-1 remained posteriorly polarized in the absence of the other Wnts tested, which may explain their functional redundancy. The posterior polarization of EGL-20 and CWN-1 could reflect amplification of their gradients, but neither the posterior polarization of CWN-2 nor its ability to polarize MOM-5 on its own can, because its gradient runs in the opposite direction. Therefore, seam cells are likely to have a polarity cue other than Wnt gradients, which may act through polarizing the receptor.

### Wnt assembles signaling components into signalosome

In our observations, Wnts and receptors, except for LIN-17/Fz, formed puncta on seam cells (Figure 1 and 3). Therefore, we analyzed whether Wnts, receptors and downstream components colocalize in puncta. We used CAM-1::mKate2(*os182*) puncta as a reference to examine its colocalization with other signaling components endogenously tagged with mNG on seam cell membrane marked with SCMp::BFP::CAAX (*osIs276*).

We observed mNG::CWN-2/Wnt (Figure 6A), EGL-20/Wnt::mNG (Figure 6B), and MOM-5/Fz::mNG (Figure 6C) puncta colocalizing with CAM-1::mKate2 on the seam cell membrane. Dishevelled (Dvl) is a core Wnt signaling component and localizes to the membrane in puncta^29^. We have shown that three Dvl homologs (DSH-1/Dvl, DSH-2/Dvl, and MIG-5/Dvl) redundantly regulate seam cell polarity, as assessed by cell fates^17^. Therefore, we examined mNG::MIG-5(*os234*; this study) and found its colocalization with CAM-1::mKate2 puncta (Figure 6I). These results indicate that Wnt, Ror, Fz, and Dvl colocalize in the same puncta and form a signalosome-like signaling complex.

**Figure 6.**
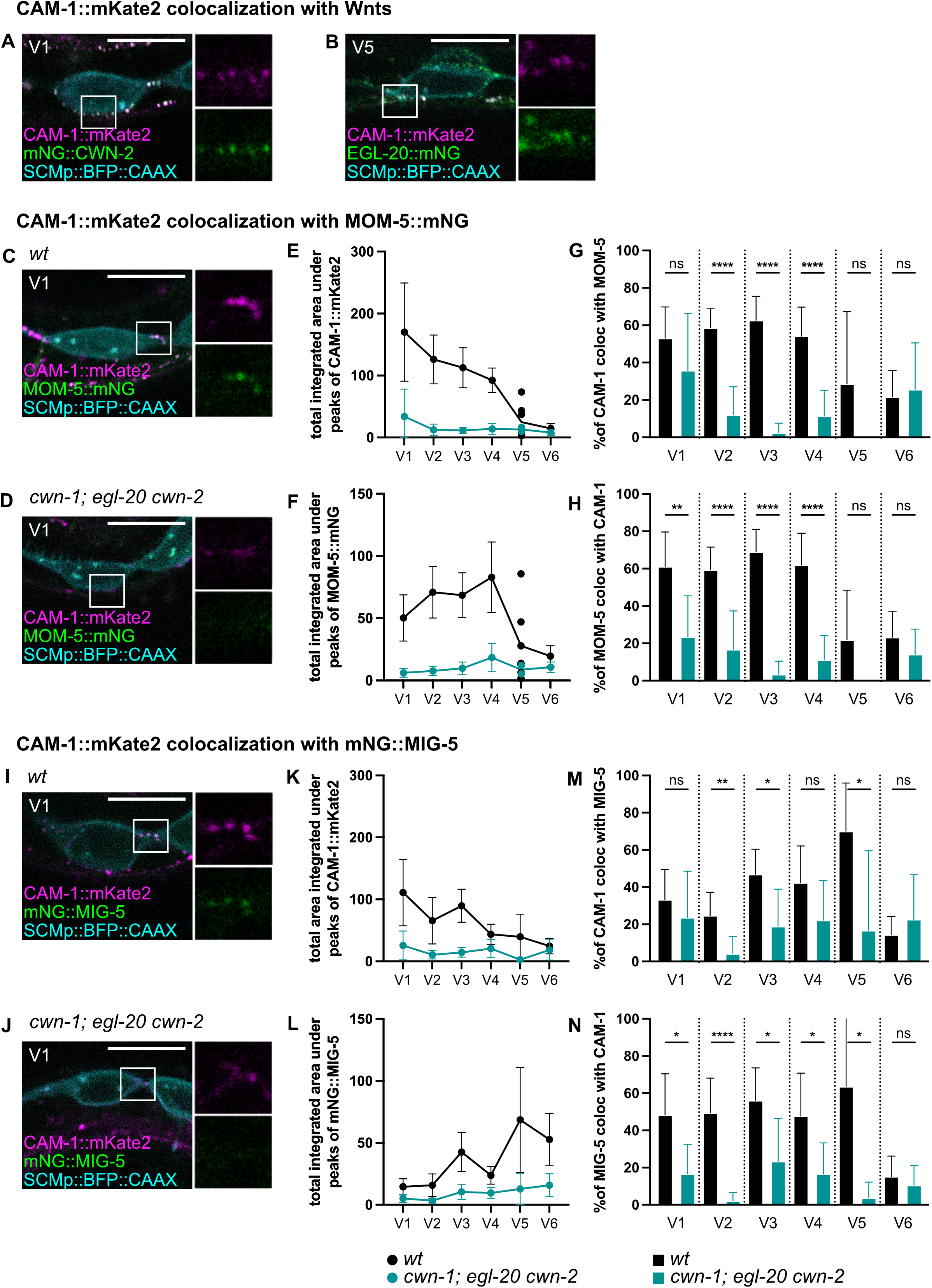
Wnts are required for the colocalization of CAM-1 with MOM-5 and MIG-5 at the seam cell membrane. **(A, B)** Representative single-plane confocal images of endogenous CAM-1::mKate2 (magenta) with mNG::CWN-2 in V1 (A) or EGL-20::mNG in V5 (B) (green), together with the seam cell membrane marker SCMp::BFP::CAAX (cyan). Boxed regions are shown magnified on the right as separate mKate2 (top) and mNG (bottom) channels. **(C, D)** Representative images of CAM-1::mKate2 (magenta), MOM-5::mNG (green), and SCMp::BFP::CAAX (cyan) in V1 of wild-type (C) and *cwn-1; egl-20 cwn-2* triple Wnt mutant (D) larvae. Boxed regions are shown magnified on the right as separate channels. **(E, F)** Total integrated area under the peaks of CAM-1::mKate2 (E) and MOM-5::mNG (F) in line profiles along the membrane of seam cells V1–V6 in wild type (black) and *cwn-1; egl-20 cwn-2* mutants (teal). **(G, H)** Percentage of CAM-1::mKate2 peaks colocalizing with MOM-5::mNG (G) and of MOM-5::mNG peaks colocalizing with CAM-1::mKate2 (H) in each seam cell in wild type (black) and *cwn-1; egl-20 cwn-2* mutants (teal). A peak was scored as colocalized when its full width at half maximum (FWHM) overlapped the FWHM footprint of a peak in the other channel (see Figure S2). Missing bars indicate 0%. **(I–N)** Same analyses as (C–H) for CAM-1::mKate2 and endogenous mNG::MIG-5: representative images in V1 of wild type (I) and *cwn-1; egl-20 cwn-2* mutants (J); total integrated area under the peaks of CAM-1::mKate2 (K) and mNG::MIG-5 (L); percentage of CAM-1 peaks colocalizing with MIG-5 (M) and of MIG-5 peaks colocalizing with CAM-1 (N). Animals were imaged at early L1 (∼3 h post-hatching). Anterior is to the left. Scale bars, 10 µm. Because the receptors are also expressed in the muscle adjacent to the ventral side of the seam cells, line profiles were drawn along the dorsal membrane only, and membranes overlapping the Q cell were excluded. Data are mean with 95% confidence intervals; n > 15 cells per seam cell position, except V4 and V5, where exclusion of Q-overlapping membranes left fewer than 10 cells in some conditions and individual data points are shown. Statistical significance was determined by t-test: ns, not significant; *P < 0.05; **P < 0.01; ***P < 0.001; ****P < 0.0001.

Next, we tested how these puncta are affected in the absence of three Wnts. Puncta in the red and green channels were identified from intensity profiles along the seam cell membrane marked with SCMp::BFP::CAAX and analyzed for their colocalization (see Methods, Figure S2). In triple Wnt mutants (*cwn-1; egl-20 cwn-2. egl-20 cwn-2* are homozygous from heterozygous mother), CAM-1::mKate2 puncta were still observed on the seam cell membrane, whereas MOM-5::mNG signal disappeared from CAM-1 puncta (Figure 6D). Quantification showed that the total integrated area under the peaks representing amounts of proteins in puncta decreased for both CAM-1 (Figure 6E) and MOM-5 (Figure 6F), and their colocalization was markedly reduced (Figure 6G-6H).

Similarly, mNG::MIG-5 signal disappeared from CAM-1 puncta in triple Wnt mutants (Figure 6J). The total integrated area under the peaks of MIG-5 also decreased (Figure 6L), together with reduced colocalization between MIG-5 and CAM-1 (Figure 6M-6N).

The results suggest that receptors and Dvl assemble into the signalosome upon Wnt signal reception, whereas in the absence of Wnts, Ror, Fz and Dvl colocalize less efficiently. Together, these results suggest that Wnts promote the assembly of receptors and Dvl into the signalosome, which is likely to underlie the reciprocal regulation of Wnt and receptor localization.

### Wnts and Dishevelled regulate each other’s localization and polarization

Since MIG-5/Dvl association into signalosome is Wnt-dependent (Figure 6I-6N), we next asked whether Wnts also regulate MIG-5 polarity. In *egl-20/*Wnt mutants, the membrane signal of mNG::MIG-5(*cp385*^26^) was comparable to that in wild-type animals, whereas its polarity on V5 was reversed (Figure 7A), consistent with the reversal of V5 polarity in *egl-20* mutants^30^. In triple Wnt mutants, MIG-5 polarity was lost in V2 and V5, and its signal decreased compared to wild-type in V1-V3 and V6 (Figure 7B and Table S4), suggesting that Wnts are required for membrane recruitment of Dvl, as shown in other organisms^31–34^. Thus, Wnts regulate the membrane localization of Dvl as well as its polarity.

**Figure 7.**
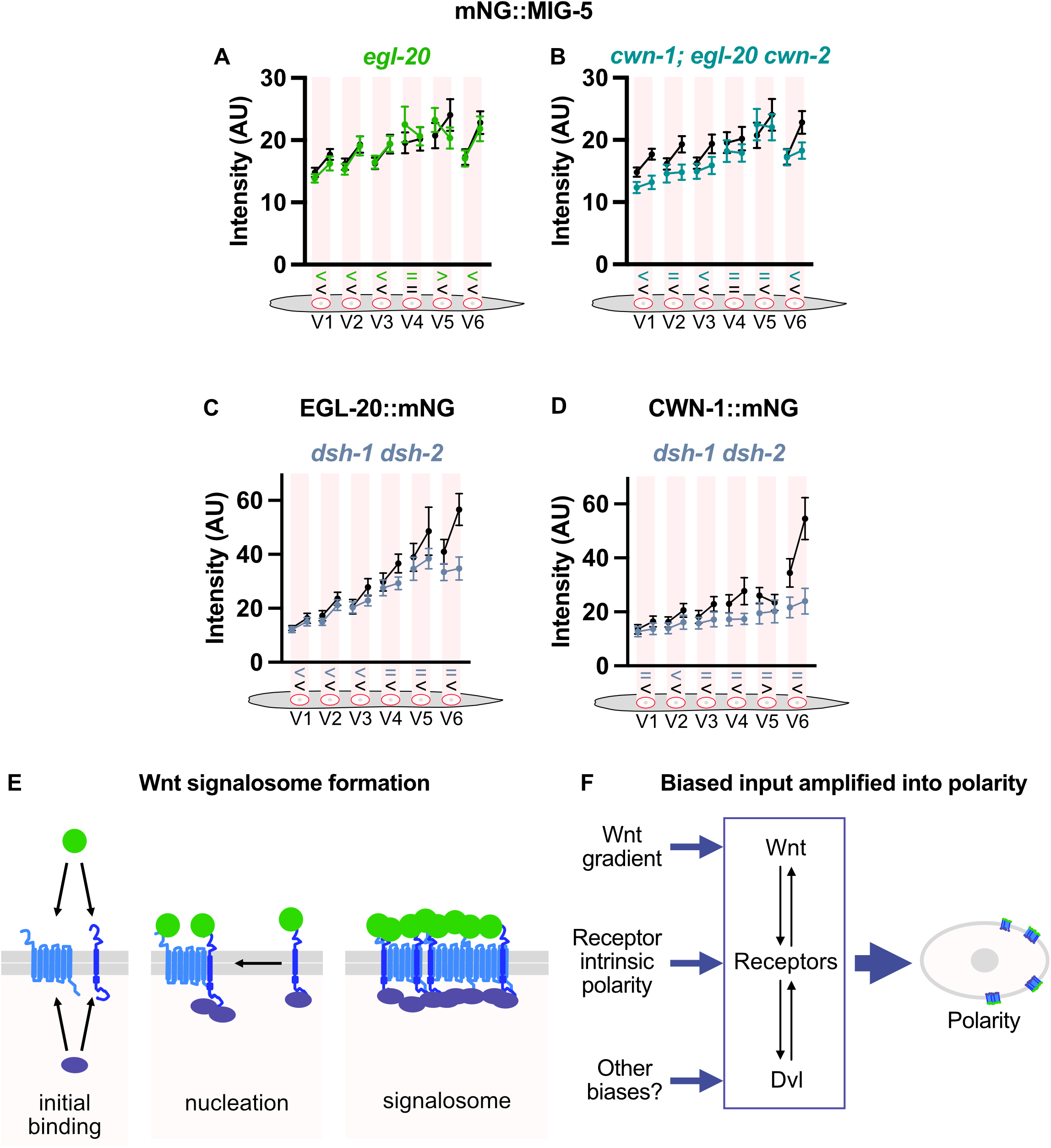
Wnts and Dishevelled promote each other’s localization and polarization through positive feedback. **(A–D)** Mean fluorescence intensity in the anterior and posterior halves of the membrane of seam cells V1–V6 for mNG::MIG-5 in *egl-20* mutants (A), in *cwn-1; egl-20 cwn-2* triple mutants (B), and for EGL-20::mNG (C) and CWN-1::mNG (D) in *dsh-1 dsh-2* double mutants (colored), compared with wild type (black). *dsh-1 dsh-2* mutants are homozygous progeny of heterozygous mothers. Within each shaded box, the left and right points are the anterior and posterior halves of the same cell. Symbols below each cell show the result of the A–P comparison (paired t-test) for the mutant (colored, top) and wild type (black, bottom) as shown in Figure 1. Animals were imaged at early L1 (∼3 h post-hatching). Intensities are raw, without background subtraction. Error bars, 95% confidence intervals. Means, SD, n, and P-values are provided in Supplemental File 1. **(E)** Model of Wnt signalosome formation. *Initial binding:* Wnt ligand (green) binds extracellularly to Frizzled (light blue) and Ror (dark blue), while Dvl (purple) binds the receptor intracellularly. *Nucleation:* A ligand–receptor–Dvl complex recruits additional Wnt-bound receptors laterally within the membrane (arrow). *Signalosome:* Further recruitment produces a higher-order cluster of Wnt, receptors and Dvl. **(F)** Model for amplification of biased inputs into cell polarity. Wnt, receptors and Dvl reinforce one another’s localization (double arrows), so that signalosome assembly is self-reinforcing. Asymmetric inputs, such as the extracellular Wnt gradient, intrinsic polarity of the receptors, and possibly other unidentified biases, determine where in the cell signalosomes preferentially nucleate and are stabilized. Positive feedback within the complex amplifies this initial bias into robust anterior–posterior polarity.

We showed that Wnts and receptors reciprocally regulate each other’s localization and polarization. We therefore asked whether Dvl, conversely, regulates the localization of Wnts. Although Dvl is classically considered a downstream component of Wnt signaling, we found that it regulates Wnt localization and polarization on the seam cell membrane. In *dsh-1/*Dvl*(or302) dsh-2/*Dvl*(ok1445)* double mutants (homozygous progeny derived from heterozygous mothers), CWN-1/Wnt::mNG became symmetric on nearly all seam cells (Figure 7D), and EGL-20::mNG polarity was lost on V4–V6 (Figure 7C). The normal asymmetry in V1–V3 is likely due to the remaining MIG-5/Dvl in *dsh-1 dsh-2* double mutants. Loss of *dsh-1* and *dsh-2* also decreased Wnt signal intensity on the membrane (Figure 7C-7D, Table S4), indicating that Dvl regulates not only the polarization of Wnts but also their membrane localization.

Together, these results show a positive feedback regulation between Wnts and Dvl. Since Wnts and Dvl are unlikely to interact directly, their interaction should occur through receptors. Therefore, our results suggest double positive-feedback loops that drive the formation of a signalosome: Wnt⇄receptor and receptor⇄Dvl. (Figure 7E-7F).

## Discussion

### Feedback regulation of Wnt localization

Wnts are well known to form graded distribution. We showed that, in. *C. elegans*, all three Wnts are globally distributed in gradients along the A–P axis (Figure 1) consistent with their sites of expression^20^. However, we showed that localization of Wnts on seam cells did not simply mirror global gradients. In particular, localization of CWN-2/Wnt was observed in the posterior side of seam cells against its global gradient (Figure 1I). Therefore, individual seam cells reorganize Wnt ligands on their membranes. Reciprocal interactions among Wnt ligands, their receptors and Dvl are likely to organize a polarized, membrane-associated signalosome (Figure 7E-7F). An important feature of such a feedback-based system is its ability to amplify spatial biases. An initial bias in any component of the feedback loops could promote the biased nucleation of the signalosome (Figure 7F). Mutual stabilization and recruitment of additional components would then amplify this bias into robust cortical polarization. Because multiple biases can act at the same time, this mechanism would allow seam cells to integrate several spatial inputs, which may explain the functional redundancy among Wnts and their receptors. It has been shown that Wnt puncta recover at the same sites after photobleaching^22^, suggesting that nucleation sites are stable. Thus, dynamic signalosome assembly may not only amplify the initial spatial bias but also help maintain the resulting spatial distribution.

The gradients of the posterior Wnts are likely candidates for the initial spatial bias, since posterior Wnts have an instructive function in the *lin-17*/Fz mutant background^17^. This hypothesis could not be tested directly, since ectopically expressed CWN-1/Wnt and EGL-20/Wnt from reversed sources are poorly localized on seam cells as described in Yamamoto et al.^21^ (and data not shown). The gradient of CWN-2 cannot be the initial bias, since its localization is opposite to the direction of its gradient. In fact, CWN-2 acts permissively in seam cell polarity, even in the *lin-17* mutant background^17^. The results indicate that the seam cell itself, rather than the CWN-2 gradient, determines where CWN-2 is localized. This may be explained by the strong dependence of CWN-2 membrane localization on CAM-1/Ror (Figure 2J, 2N and S1B), which retains posterior polarity in anterior seam cells even in the absence of all three Wnts (Figure 4E). We therefore speculate that the intrinsic polarity of CAM-1 provides a spatial bias that overrides the CWN-2 gradient. A similar mechanism may also underlie asymmetric localization of CWN-1 and EGL-20, consistent with their permissive functions in the absence of *lin-17* mutation^17,21^. Therefore, both Wnt gradients and intrinsic polarity represented by CAM-1 localization as well as other intracellular biases might be used as initial cues to be amplified during cell polarization, in the case of the posterior Wnts.

Our results also explain previous genetic findings that CWN-2 functions mainly through CAM-1. We showed that *cwn-1; egl-20 cwn-2* and *cam-1 cwn-1; egl-20* triple mutants exhibit similar seam cell polarity defects, and that loss of *cwn-2* does not enhance the defects of *cam-1* mutants^17^. Consistent with these genetic evidences for CAM-1 dependence of CWN-2 function, our present results show that CWN-2 fails to localize on any target cells in *cam-1* mutants (Figure 2N and S1B). This specific dependence of CWN-2 on CAM-1 may be evolutionarily conserved, because CWN-2 and CAM-1 are orthologs of Wnt5^35^ and Ror, respectively. Wnt5 and Ror are a well-established ligand–receptor pair and are considered core components of non-canonical Wnt signaling^36^.

### Comparison with migration model

Migrating cells use an intracellular amplification mechanism during polarization^37^. Our results show that seam cells use an amplification system that acts at the level of establishing cortical polarity. An interesting implication of our model is that seam cells do not simply amplify their response to Wnt signals, but also modify the signal itself. Because cells ultimately respond to receptor-bound rather than freely diffusing Wnt, controlling where Wnt is localized on the membrane allows a cell to determine its own response. In this way, seam cells may sharpen the posterior bias provided by the CWN-1/Wnt and EGL-20/Wnt gradients, and override the opposing gradient of CWN-2/Wnt. This is reminiscent of migrating cells that reshape the distribution of their guidance cues, such as the zebrafish lateral line primordium. There, the decoy receptor Cxcr7 sequesters the chemokine Cxcl12a at the rear of the tissue, generating a gradient across the migrating primordium^7,8^. In contrast to migrating cells, which modify the distribution of a free ligand in their surroundings, seam cells modify ligand distribution locally, by controlling where Wnt is retained on their own membranes.

In addition to this amplification of the signal on the cell membrane, a second layer of amplification is likely to act on the response, because signalosomes, although enriched on the posterior side, are also present on the anterior side, whereas the resulting cell fate decision is binary. Consistently, as we mentioned earlier, in *cam-1/*Ror mutants, none of the three Wnts showed significant asymmetry in V1 (Figure 2A, 2F and 2J), yet V1 daughter cell fates were still correctly polarized in the majority of animals^17,38^, indicating that even weak cortical asymmetry might be amplified to produce a proper binary outcome.

### Signalosome assembly in Wnt signaling

In both non-canonical and canonical Wnt signaling, membrane-associated signaling complexes containing Wnt, receptors and Dishevelled (Dvl) proteins play essential roles. In PCP signaling, core PCP proteins assemble into asymmetric, signalosome-like complexes at cell boundaries to establish and coordinate cell polarity^12^. Dvl also promotes the clustering and stabilization of PCP complexes, with its DIX domain contributing to Dvl3 asymmetric localization^39^. In canonical Wnt signaling, Wnt promotes the formation of receptor-associated signaling complexes, or signalosomes, through multivalent interactions with its receptors^13–15^, as well as Dvl polymerization mediated by its DIX and DEP domains^16^. ROR2 can also participate in receptor complexes, cooperating with FZD7 to promote WNT5A-induced Dvl polymerization^40^ and associating with FZD8- and LRP6-containing signalosomes^41^. However, whether receptor–Dvl-containing complexes contribute to the spatial distribution of Wnt ligands remains unclear. It has been recently shown that the core component Vangl regulates the polarized localization of Wnt^42^. However, to our knowledge, our results provide the first evidence that Dvl regulates the localization of Wnt ligands themselves. Moreover, we showed that asymmetric seam cell divisions and polarized EGL-20/Wnt localization do not involve Vangl. Therefore, our results, for the first time, show Vangl-independent regulation of Wnt localization by the receptors and Dvl. Although CAM-1/Ror is involved in Wnt localization in *C. elegans*, asymmetric localization of EGL-20 in V3-V6 cells in *cam-1* mutants (Figure 2A) raises a possibility that the feedback regulation might occur without Ror through Wnt-Fz-Dvl interactions even in canonical Wnt signaling.

## Materials and Methods

### Worm Culture

N2 Bristol was used as the wild-type strain. The animals were cultured at 22.5°C on NGM plate seeded with *E. coli* OP50.

### Generation of strains

Full strain and allele list with references is given in Supplemental File 1.

*osIs33 [SCMp::mCherry::CAAX] and osIs276 [SCMp::BFP::CAAX] –* multicopy UV integration of extrachromosomal arrays, obtained by injection of pYM33 or pMA110 with pUNC76 into *unc-76 (e911)* background.

*osIs114* and *osIs118 [eft-3p::TIR-1 + coelRFP]* were generated by multicopy UV integration of pMA25 with pUNC76 into *unc-76 (e911)* background.

*osIs253 [Peft-3::TIR1(F79G)::mRuby::tbb-2 3’UTR] and osIs182 [eft-3::TIR1(F79G) +* Cre-excisable selection cassette carrying *myo-3p::GFP* and a neomycin-resistance gene] were generated by CRISPR/Cas9-mediated insertion at the *oxTi365* site on chromosome V, using the self-excising cassette method^43^. Repair templates were based on pOB5, and pTK73_pOB5 was used for sgRNA expression^44^.

*lin-17(os229)* [*lin-17::mNG::3xFLAG*] was generated by CRISPR/Cas9-mediated insertion using the self-excising cassette method^43^, with a repair template based on pDD268 (Addgene #132523). The tag was inserted immediately before the *lin-17* stop codon (sgRNA 5′-TCTCGCTCAGACGACCTTAC-3′). *mig-5(os234)* [*mNG::3xMyc::mig-5*] was generated in the same way in the *os182* background, at the same site as *cp385*, using pDD162 as the sgRNA vector as described in Heppert et al.^26^. Its repair template was based on pDD287 (Addgene #70685), with mKate2 replaced by mNG from pDD268. For both alleles, edited animals were selected by hygromycin resistance and the Rol phenotype, and the cassette was excised by heat-shock-induced Cre.

*cwn-1(cp205)* [*cwn-1::mNG::3xFLAG*] was obtained from the same injection round as *cp209*, described in Pani et al.^23^.

*cam-1(os182)* [*cam-1::mAID::mKate2*] was generated by CRISPR/Cas9-mediated homologous recombination. The repair template carried mAID^45,46^ and mKate2 flanked by 120-bp homology arms corresponding to the regions immediately upstream and downstream of the *cam-1* stop codon. A crRNA (5′-AGAGGATGGTGATTCTGATT-3′) was duplexed with Alt-R tracrRNA (IDT), complexed with Alt-R *S.p.* Cas9 nuclease V3 (IDT), and injected into N2 together with the repair template and pRF4 [*rol-6(su1006)*]. Insertion was confirmed by PCR and sequencing.

*egl-20(os214)* was generated by CRISPR/Cas9-mediated non-homologous end-joining repair in the *cwn-2(os187); osIs33* background. Two crRNAs targeting *egl-20* (5′-CGTTTATGGGCAATATTCTC-3′ and 5′-CGCAACAAGTTCCATATAAC-3′) were duplexed with Alt-R CRISPR-Cas9 tracrRNA, complexed with Alt-R *S.p.* Cas9 nuclease V3 (IDT), and co-injected with a *dpy-10* crRNA. Error-prone end-joining repair yielded a 2-bp deletion (c.99_100del) that introduces a premature stop codon (p.(Tyr33Ter)).

### Microscopy and imaging

For *in vivo* imaging, worms were immobilized on 4% agar pads containing 0.1 M sodium azide. Imaging was performed on a Zeiss LSM700 confocal microscope with a 63×, N.A. 1.40 oil-immersion objective. All animals were imaged at the L1 stage, after the initial Q-cell migration and before seam cell division, which corresponds to around 3 hours after hatching.

Each mutant dataset was compared with control animals imaged on the same microscope using identical acquisition settings. To avoid the effect of laser power degradation, control and mutant data were acquired within one year of each other.

### AID degron experiments

Strains carrying *osIs114* or *osIs118* were treated with indole-3-acetic acid (IAA), and strains carrying *osIs253 and osIs182* with 5-phenyl-indole-3-acetic acid (5-Ph-IAA), each dissolved in dimethyl sulfoxide (DMSO). Eggs were hatched on plates containing 10 µM IAA or 5-Ph-IAA, and were imaged after the initial Q-cell migration and before seam cell divisions, which corresponds to around 3 hours after hatching.

### Quantification of Polarity

For Wnt and Dvl dorsal and ventral side of each seam cell was hand-traced with 4px thick spine-fit segmented line using Fiji in anterior-to-posterior direction, using *osIs33* signal as a guideline. For receptors only the dorsal side was quantified because of the presence of the signal in the ventral muscle, and receptor signal cannot be differentiated whether it belongs to the seam cell or muscle. For receptors and Dvl V4 and V5 cells contacting Q cell and its daughters were excluded for the same reason. One-dimensional intensity plot was extracted for each line, midpoint was found for each dorsal and ventral line, and anterior and posterior was averaged for each cell, with the account for the length. To compare anterior and posterior of each cell, paired t-test was used.

Fates of seam cell daughters in the *vang-1* mutant was analyzed by the *elt-3p*::GFP (*wgIs559*) expression in hyp7 in the late L1 stage.

### Quantification of Colocalization

The dorsal membrane was traced in each animal using the *osIs276* signal as a guide, and fluorescence intensity along the trace was extracted as a one-dimensional profile for each channel (pixel size 0.062 µm). Baseline was defined as the median intensity, and noise as σ = 1.4826 × MAD (median absolute deviation). Peaks were accepted if they exceeded baseline by ≥3σ, had a prominence of ≥3σ, and were ≥6 pixels (0.37 µm) from neighboring peaks. They also had to be at least as wide at half-height as the confocal diffraction limit (0.4λ/NA; NA 1.4; 0.145 µm green, 0.171 µm red). Each peak’s footprint was the interval above half-maximum relative to baseline. Integrated intensity was the baseline-subtracted area under the peak.

Colocalization was scored by footprint overlap: a peak was counted as colocalized if its footprint overlapped any peak in the other channel (Jaccard index > 0). This was calculated in both directions (% red overlapping green, % green overlapping red). Analyses were run in MATLAB (Signal Processing Toolbox). The integrated area under the peaks above the baseline was calculated summarizing all the peaks on the same cell. A representative example of the method is given in Figure S2.

## Acknowledgement

We would like to thank Katherine Long for comments on the manuscript. Some strains were provided by the CGC, which is funded by NIH Office of Research Infrastructure Programs (P40 OD010440), and by the National Bioresource Project of Japan (NBRP). This research was supported by JSPS KAKENHI grants (JP16KT0078, JP23K21314 to HS and JP23K05836 to TN) and the Takeda Science Foundation, the Ohsumi Frontier Science Foundation to HS, and NIH grant R35GM134838 to BG and R35GM142880 to AP.

---

**Figure S1. CWN-2 accumulation on target cells depends on CAM-1. (A)** Representative single-plane confocal images of endogenous mNG::CWN-2 in the head region of a wild-type early L1 larva (∼3 h post-hatching), showing a mid-plane (top) and a surface plane (bottom) of the same animal. mNG::CWN-2 is detected in the producing cells (pharynx, nerve ring, and ventral cord) and on target cells such as seam cell V1. Coelomocytes, which take up secreted proteins from the body cavity, are also labeled.

**(B)** Representative single-plane confocal image (surface plane) of mNG::CWN-2 in a *cam-1* early L1 larva. mNG::CWN-2 is lost from target cells such as V1 but remains detectable in producing cells and coelomocytes. Anterior is to the left. Scale bars, 10 µm.

**Figure S2. Example of line-profile analysis used to quantify colocalization.** Representative fluorescence intensity profiles of CAM-1::mKate2 (magenta) and MOM-5::mNG (green) along the dorsal membrane of seam cell V1 in a wild-type early L1 larva (sample #3). The dashed line indicates the baseline (median intensity of the profile), and the dotted line indicates the peak detection threshold, set at 3σ above the baseline (σ = 1.4826 × median absolute deviation of the profile). Peaks exceeding the threshold are shaded, and the shaded area above the baseline corresponds to the integrated area under the peaks quantified in Figure 6E, F, K, L. Horizontal bars mark the full width at half maximum (FWHM) of each peak. A peak was scored as colocalized when its FWHM overlapped the FWHM of a peak in the other channel. In this example, two CAM-1 and two MOM-5 peaks were detected; one pair is colocalized and the remaining two peaks are not. Intensity is in arbitrary units.

**Table S1. Comparison of Wnt intensity between wild-type and *cam-1* mutant seam cells.** Mean fluorescence intensity along the whole membrane of each seam cell (V1–V6) for EGL-20::mNG, CWN-1::mNG, and mNG::CWN-2 in control and *cam-1(gm122)* mutant animals, corresponding to Figure 2. “Difference with control” is the control mean minus the mutant mean; positive values indicate lower intensity in the mutant. P-values are from two-tailed Welch’s t-tests comparing mutant and control. Intensities are raw, in arbitrary units.

**Table S2. Comparison of Wnt intensity between neighboring seam cells.** Mean fluorescence intensity along the whole membrane of each seam cell (V1–V6) for EGL-20::mNG in control, *cam-1*, and *mom-5(AID); cam-1* animals, and for CWN-1::mNG in control and *cam-1* animals. Control values correspond to Figure 1D, E. “Difference with next cell” is the mean intensity of each cell minus that of its posterior neighbor (e.g., V1 − V2); negative values indicate higher intensity in the more posterior cell. P-values are from paired t-tests comparing each cell with its posterior neighbor within the same animals. NA, not applicable (V6 has no posterior neighbor). Intensities are raw, in arbitrary units.

**Table S3. Comparison of receptor intensity between wild-type and Wnt mutant seam cells.** Mean fluorescence intensity along the whole membrane of each seam cell (V1–V6) for CAM-1::mNG and MOM-5::mNG in control, *cwn-1; egl-20* double mutant, and *cwn-1; egl-20 cwn-2* triple mutant animals, corresponding to Figure 4. “Difference with control” is the control mean minus the mutant mean; negative values indicate higher intensity in the mutant. P-values are from two-tailed Welch’s t-tests comparing mutant and control. For MOM-5::mNG, separate controls imaged with each mutant are shown. Intensities are raw, in arbitrary units.

**Table S4. Comparison of MIG-5 and Wnt intensity between wild-type and mutant seam cells.** Mean fluorescence intensity along the whole membrane of each seam cell (V1–V6) for mNG::MIG-5 in control and *cwn-1; egl-20 cwn-2* triple mutant animals, and for EGL-20::mNG and CWN-1::mNG in control and *dsh-1 dsh-2* double mutant animals (homozygous progeny from heterozygous mothers), corresponding to Figure 7. “Difference with control” is the control mean minus the mutant mean; positive values indicate lower intensity in the mutant. P-values are from two-tailed Welch’s t-tests comparing mutant and control. Intensities are raw, in arbitrary units.

## References

1. Brandman, O., and Meyer, T. (2008). Feedback Loops Shape Cellular Signals in Space and Time. Science 322, 390–395. 10.1126/science.1160617.

2. Sasaki, A.T., Janetopoulos, C., Lee, S., Charest, P.G., Takeda, K., Sundheimer, L.W., Meili, R., Devreotes, P.N., and Firtel, R.A. (2007). G protein–independent Ras/PI3K/F-actin circuit regulates basic cell motility. The Journal of Cell Biology 178, 185–191. 10.1083/jcb.200611138.

3. Weiner, O.D., Neilsen, P.O., Prestwich, G.D., Kirschner, M.W., Cantley, L.C., and Bourne, H.R. (2002). A PtdInsP3- and Rho GTPase-mediated positive feedback loop regulates neutrophil polarity. Nat Cell Biol 4, 509–513. 10.1038/ncb811.

4. Reichman-Fried, M., Minina, S., and Raz, E. (2004). Autonomous Modes of Behavior in Primordial Germ Cell Migration. Developmental Cell 6, 589–596. 10.1016/S1534-5807(04)00074-7.

5. Petrie Aronin, C.E., Zhao, Y.M., Yoon, J.S., Morgan, N.Y., Prüstel, T., Germain, R.N., and Meier-Schellersheim, M. (2017). Migrating Myeloid Cells Sense Temporal Dynamics of Chemoattractant Concentrations. Immunity 47, 862–874.e3. 10.1016/j.immuni.2017.10.020.

6. Georgantzoglou, A., Poplimont, H., Walker, H.A., Lämmermann, T., and Sarris, M. (2022). A two-step search and run response to gradients shapes leukocyte navigation in vivo. Journal of Cell Biology 221, e202103207. 10.1083/jcb.202103207.

7. Donà, E., Barry, J.D., Valentin, G., Quirin, C., Khmelinskii, A., Kunze, A., Durdu, S., Newton, L.R., Fernandez-Minan, A., Huber, W., et al. (2013). Directional tissue migration through a self-generated chemokine gradient. Nature 503, 285–289. 10.1038/nature12635.

8. Venkiteswaran, G., Lewellis, S.W., Wang, J., Reynolds, E., Nicholson, C., and Knaut, H. (2013). Generation and Dynamics of an Endogenous, Self-Generated Signaling Gradient across a Migrating Tissue. Cell 155, 674–687. 10.1016/j.cell.2013.09.046.

9. Tweedy, L., Thomason, P.A., Paschke, P.I., Martin, K., Machesky, L.M., Zagnoni, M., and Insall, R.H. (2020). Seeing around corners: Cells solve mazes and respond at a distance using attractant breakdown. Science 369, eaay9792. 10.1126/science.aay9792.

10. Minegishi, K., Hashimoto, M., Ajima, R., Takaoka, K., Shinohara, K., Ikawa, Y., Nishimura, H., McMahon, A.P., Willert, K., Okada, Y., et al. (2017). A Wnt5 Activity Asymmetry and Intercellular Signaling via PCP Proteins Polarize Node Cells for Left-Right Symmetry Breaking. Developmental Cell 40, 439–452.e4. 10.1016/j.devcel.2017.02.010.

11. Gao, B., Song, H., Bishop, K., Elliot, G., Garrett, L., English, M.A., Andre, P., Robinson, J., Sood, R., Minami, Y., et al. (2011). Wnt Signaling Gradients Establish Planar Cell Polarity by Inducing Vangl2 Phosphorylation through Ror2. Developmental Cell 20, 163–176. 10.1016/j.devcel.2011.01.001.

12. Strutt, H., Gamage, J., and Strutt, D. (2016). Robust Asymmetric Localization of Planar Polarity Proteins Is Associated with Organization into Signalosome-like Domains of Variable Stoichiometry. Cell Reports 17, 2660–2671. 10.1016/j.celrep.2016.11.021.

13. DeBruine, Z.J., Ke, J., Harikumar, K.G., Gu, X., Borowsky, P., Williams, B.O., Xu, W., Miller, L.J., Xu, H.E., and Melcher, K. (2017). Wnt5a promotes Frizzled-4 signalosome assembly by stabilizing cysteine-rich domain dimerization. Genes Dev. 31, 916–926. 10.1101/gad.298331.117.

14. Nile, A.H., Mukund, S., Stanger, K., Wang, W., and Hannoush, R.N. (2017). Unsaturated fatty acyl recognition by Frizzled receptors mediates dimerization upon Wnt ligand binding. Proc. Natl. Acad. Sci. U.S.A. 114, 4147–4152. 10.1073/pnas.1618293114.

15. Yue, D., Sun, G., Cao, Y., Li, H., Yang, Y., Pan, Z., Xue, L., Zhang, L., Wang, Z., and Xu, W. (2026). Structural basis of Wnt signalosome extracellular complex assembly. Cell 189, 4310–4324.e18. 10.1016/j.cell.2026.05.006.

16. Gammons, M.V., Renko, M., Johnson, C.M., Rutherford, T.J., and Bienz, M. (2016). Wnt Signalosome Assembly by DEP Domain Swapping of Dishevelled. Molecular Cell 64, 92–104. 10.1016/j.molcel.2016.08.026.

17. Sawa, H., Asakawa, M., and Negishi, T. (2026). Differential functions of multiple Wnts and receptors in cell polarity regulation in *Caenorhabditis elegans*. GENETICS 232, iyag004. 10.1093/genetics/iyag004.

18. Sawa, H., and Korswagen, H.C. (2013). Wnt signaling in C. elegans. WormBook, 1–30. 10.1895/wormbook.1.7.2.

19. Mizumoto, K., and Sawa, H. (2007). Cortical β-Catenin and APC Regulate Asymmetric Nuclear β-Catenin Localization during Asymmetric Cell Division in C. elegans. Developmental Cell 12, 287–299. 10.1016/j.devcel.2007.01.004.

20. Harterink, M., Kim, D.H., Middelkoop, T.C., Doan, T.D., Van Oudenaarden, A., and Korswagen, H.C. (2011). Neuroblast migration along the anteroposterior axis of *C. elegans* is controlled by opposing gradients of Wnts and a secreted Frizzled-related protein. Development 138, 2915–2924. 10.1242/dev.064733.

21. Yamamoto, Y., Takeshita, H., and Sawa, H. (2011). Multiple Wnts Redundantly Control Polarity Orientation in Caenorhabditis elegans Epithelial Stem Cells. PLoS Genet 7, e1002308. 10.1371/journal.pgen.1002308.

22. Pani, A.M., and Goldstein, B. (2018). Direct visualization of a native Wnt in vivo reveals that a long-range Wnt gradient forms by extracellular dispersal. eLife 7, e38325. 10.7554/eLife.38325.

23. Pani, A.M., Favichia, M., and Goldstein, B. (2023). Long-distance Wnt transport in axons highlights cell type-specific modes of Wnt transport *in vivo*. Preprint at bioRxiv, 10.1101/2023.05.03.539245 https://doi.org/10.1101/2023.05.03.539245.

24. Park, F.D., Tenlen, J.R., and Priess, J.R. (2004). C. elegans MOM-5/Frizzled Functions in MOM-2/Wnt-Independent Cell Polarity and Is Localized Asymmetrically prior to Cell Division. Current Biology 14, 2252–2258. 10.1016/j.cub.2004.12.019.

25. Goldstein, B., Takeshita, H., Mizumoto, K., and Sawa, H. (2006). Wnt Signals Can Function as Positional Cues in Establishing Cell Polarity. Developmental Cell 10, 391–396. 10.1016/j.devcel.2005.12.016.

26. Heppert, J.K., Pani, A.M., Roberts, A.M., Dickinson, D.J., and Goldstein, B. (2018). A CRISPR Tagging-Based Screen Reveals Localized Players in Wnt-Directed Asymmetric Cell Division. Genetics 208, 1147–1164. 10.1534/genetics.117.300487.

27. Ferrando-Marco, M., Berger, S., and Barkoulas, M. (2026). Transcriptional feedback targeting Wnt pathway components reveals hidden heterogeneity in *Caenorhabditis elegans* seam cell lineages. GENETICS 233, iyag147. 10.1093/genetics/iyag147.

28. Ji, N., Middelkoop, T.C., Mentink, R.A., Betist, M.C., Tonegawa, S., Mooijman, D., Korswagen, H.C., and van Oudenaarden, A. (2013). Feedback Control of Gene Expression Variability in the Caenorhabditis elegans Wnt Pathway. Cell 155, 869–880. 10.1016/j.cell.2013.09.060.

29. Bienz, M. (2014). Signalosome assembly by domains undergoing dynamic head-to-tail polymerization. Trends in Biochemical Sciences 39, 487–495. 10.1016/j.tibs.2014.08.006.

30. Whangbo, J., Harris, J., and Kenyon, C. (2000). Multiple levels of regulation specify the polarity of an asymmetric cell division in *C. elegans*. Development 127, 4587–4598. 10.1242/dev.127.21.4587.

31. Witzel, S., Zimyanin, V., Carreira-Barbosa, F., Tada, M., and Heisenberg, C.-P. (2006). Wnt11 controls cell contact persistence by local accumulation of Frizzled 7 at the plasma membrane. The Journal of Cell Biology 175, 791–802. 10.1083/jcb.200606017.

32. Bilić, J., Huang, Y.-L., Davidson, G., Zimmermann, T., Cruciat, C.-M., Bienz, M., and Niehrs, C. (2007). Wnt Induces LRP6 Signalosomes and Promotes Dishevelled-Dependent LRP6 Phosphorylation. Science 316, 1619–1622. 10.1126/science.1137065.

33. Yamanaka, H., and Nishida, E. (2007). Wnt11 stimulation induces polarized accumulation of Dishevelled at apical adherens junctions through Frizzled7. Genes to Cells 12, 961–967. 10.1111/j.1365-2443.2007.01106.x.

34. Ma, W., Chen, M., Kang, H., Steinhart, Z., Angers, S., He, X., and Kirschner, M.W. (2020). Single-molecule dynamics of Dishevelled at the plasma membrane and Wnt pathway activation. Proc. Natl. Acad. Sci. U.S.A. 117, 16690–16701. 10.1073/pnas.1910547117.

35. Prud’homme, B., Lartillot, N., Balavoine, G., Adoutte, A., and Vervoort, M. (2002). Phylogenetic Analysis of the Wnt Gene Family. Current Biology 12, 1395– 1400. 10.1016/S0960-9822(02)01068-0.

36. Konopelski Snavely, S.E., Srinivasan, S., Dreyer, C.A., Tan, J., Carraway, K.L., and Ho, H.-Y.H. (2023). Non-canonical WNT5A-ROR signaling: New perspectives on an ancient developmental pathway. In Current Topics in Developmental Biology (Elsevier), pp. 195–227. 10.1016/bs.ctdb.2023.01.009.

37. Kölsch, V., Charest, P.G., and Firtel, R.A. (2008). The regulation of cell motility and chemotaxis by phospholipid signaling. Journal of Cell Science 121, 551–559. 10.1242/jcs.023333.

38. Forrester, W.C., Dell, M., Perens, E., and Garriga, G. (1999). A C. elegans Ror receptor tyrosine kinase regulates cell motility and asymmetric cell division. Nature 400, 881–885. 10.1038/23722.

39. Sil, P., Trejo, B., Little, K.A., and Devenport, D. (2026). Dishevelled-mediated clustering stabilizes Frizzled6 and Vangl2 to establish planar cell polarity in the mammalian skin. Sci. Adv. 12, eaeh6920. 10.1126/sciadv.aeh6920.

40. Nishita, M., Itsukushima, S., Nomachi, A., Endo, M., Wang, Z., Inaba, D., Qiao, S., Takada, S., Kikuchi, A., and Minami, Y. (2010). Ror2/Frizzled Complex Mediates Wnt5a-Induced AP-1 Activation by Regulating Dishevelled Polymerization. Molecular and Cellular Biology 30, 3610–3619. 10.1128/MCB.00177-10.

41. Philippi, M., Dohle, J., Watrinet, I., Holtmannspötter, M., Miao, Y., Li, J., Birkholz, O., Rothbauer, U., Garcia, K.C., Kurre, R., et al. (2026). Hour-scale single-molecule imaging reveals dynamic assembly of the Wnt co-receptors LRP6 and ROR2 into common signalosomes. Sci. Signal. 19, eadr8141. 10.1126/scisignal.adr8141.

42. Mii, Y., Suzuki, M., Koyama, H., Nakayama, K., Takada, R., Kobayashi, T., Tsutsumi, M., Nemoto, T., Matsuyama, M., Fujimori, T., et al. (2026). Planar cell polarity emerges through polarized accumulation of Wnt11. Sci. Adv. 12, eaea0326. 10.1126/sciadv.aea0326.

43. Dickinson, D.J., Pani, A.M., Heppert, J.K., Higgins, C.D., and Goldstein, B. (2015). Streamlined Genome Engineering with a Self-Excising Drug Selection Cassette. Genetics 200, 1035–1049. 10.1534/genetics.115.178335.

44. Obinata, H., Sugimoto, A., and Niwa, S. (2018). Streptothricin acetyl transferase 2 (Sat2): A dominant selection marker for Caenorhabditis elegans genome editing. PLoS ONE 13, e0197128. 10.1371/journal.pone.0197128.

45. Natsume, T., Kiyomitsu, T., Saga, Y., and Kanemaki, M.T. (2016). Rapid Protein Depletion in Human Cells by Auxin-Inducible Degron Tagging with Short Homology Donors. Cell Reports 15, 210–218. 10.1016/j.celrep.2016.03.001.

46. Negishi, T., Asakawa, M., Kanemaki, M., and Sawa, H. (2019). Modified auxin improves the auxin-inducible degradation (AID) system for laid C. elegans embryos. MicroPubl Biol 2019. 10.17912/micropub.biology.000190.

