## supplemental Figures and Tables for "Reciprocal regulation of Wnt, receptor and Dvl localizations during cell polarization"

Figure S1

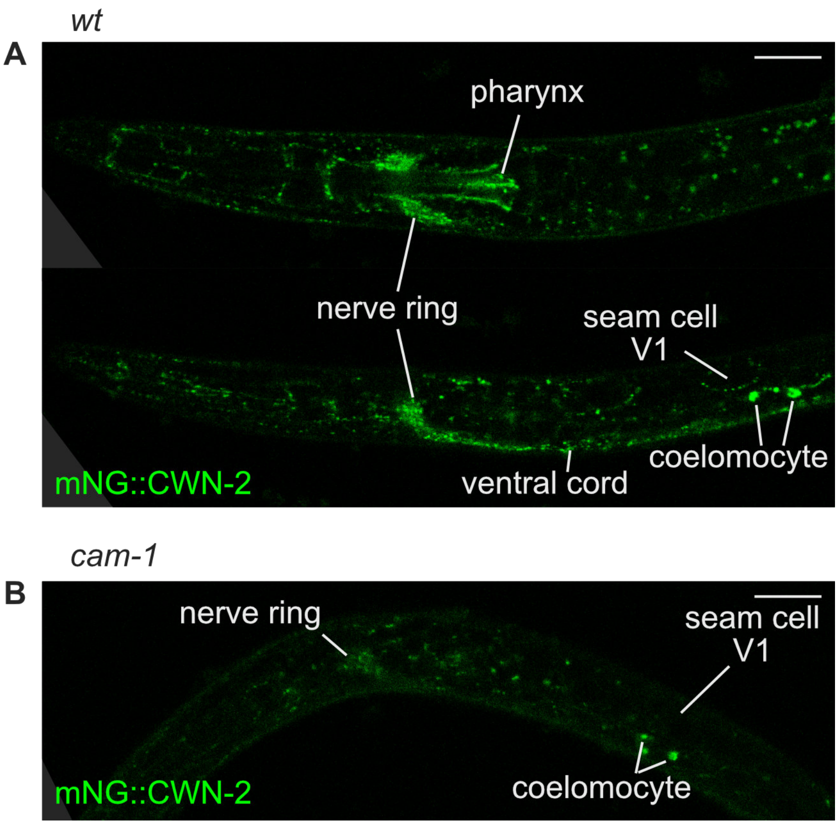
wt
A
pharynx
nerve ring
seam cell
V1
coelomocyte
ventral cord
mNG::CWN-2
cam-1
B
nerve ring
seam cell
V1
coelomocyte
mNG::CWN-2

**Figure S2**

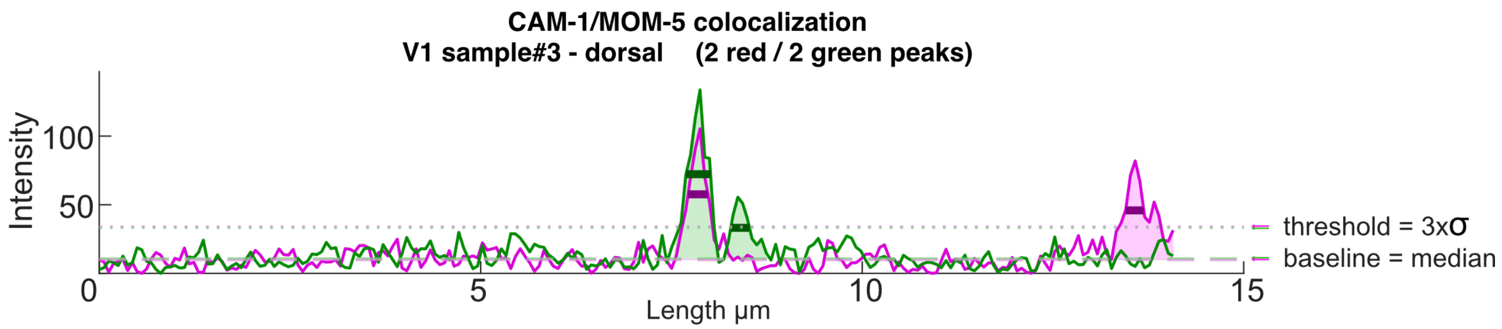
CAM-1/MOM-5 colocalization
V1 sample#3 - dorsal (2 red / 2 green peaks)
Intensity
100
50
0
5
10
15
Length µm
threshold = 3xσ
baseline = median

**Table S1**

| **EGL-20::mNG intensity average per cell** | | | | | | |
|---|---|---|---|---|---|---|
| | V1 | V2 | V3 | V4 | V5 | V6 |
| ***control*** | **14.512751** | **20.379104** | **24.212777** | **33.215453** | **43.732358** | **48.812658** |
| ***cam-1*** | **12.432187** | **13.961316** | **16.292624** | **21.561990** | **32.620540** | **33.306367** |
| difference with control | 2.080565 | 6.417788 | 7.920153 | 11.653463 | 11.111818 | 15.506291 |
| t-test p-value | 0.005940 | 1.07245 E-06 | 1.09869 E-06 | 9.7161 E-09 | 0.002816 | 8.81934 E-07 |
| **CWN-1::mNG intensity average per cell** | | | | | | |
| | V1 | V2 | V3 | V4 | V5 | V6 |
| ***control*** | **15.006468** | **18.397206** | **20.516777** | **25.345054** | **24.754023** | **44.508472** |
| ***cam-1*** | **11.212148** | **13.090992** | **14.352518** | **16.982342** | **19.532900** | **30.200781** |
| difference with control | 3.794320 | 5.306215 | 6.164258 | 8.362712 | 5.221123 | 14.307691 |
| t-test p-value | 0.000203 | 1.10987 E-05 | 1.55558 E-05 | 0.000335 | 0.003254 | 0.000161 |
| **mNG::CWN-2 intensity average per cell** | | | | | | |
| | V1 | V2 | V3 | V4 | V5 | V6 |
| ***control*** | **29.610368** | **23.670840** | **17.052952** | **16.487783** | **14.173964** | **13.374306** |
| ***cam-1*** | **15.286800** | **16.278516** | **17.121194** | **17.103160** | **15.473418** | **14.813684** |
| difference with control | 14.323568 | 7.392324 | -0.068241 | -0.615377 | -1.299454 | -1.439378 |
| t-test p-value | 1.31691 E-14 | 1.58271 E-09 | 0.916933 | 0.204979 | 0.009030 | 0.001924 |

**Table S2**

| | | value | V1 | V2 | V3 | V4 | V5 | V6 |
|---|---|---|---|---|---|---|---|---|
| EGL-20::mNG | *control* | **mean intensity** | **14.512751** | **20.379104** | **24.212777** | **33.215453** | **43.732358** | **48.812658** |
| | | difference with next cell | -5.866353 | -3.833673 | -9.002676 | -10.516905 | -5.080300 | NA |
| | | paired t-test p-value | 5.82042E-09 | 0.000253 | 1.96114E-06 | 0.001052 | 0.150337 | NA |
| | *cam-1* | **mean intensity** | **12.432187** | **13.961316** | **16.292624** | **21.561990** | **32.620540** | **33.306367** |
| | | difference with next cell | -1.529129 | -2.331308 | -5.269366 | -11.058551 | -0.685827 | NA |
| | | paired t-test p-value | 0.000150 | 3.61322E-07 | 1.77349E-08 | 1.86551E-07 | 0.730396 | NA |
| | *mom-5(AID); cam-1* | **mean intensity** | **12.109872** | **13.507310** | **15.377476** | **19.299976** | **24.500428** | **34.416858** |
| | | difference with next cell | -1.397437 | -1.870166 | -3.922500 | -5.200452 | -9.916430 | NA |
| | | paired t-test p-value | 0.000350 | 5.73558E-06 | 8.7854E-08 | 0.000251 | 1.00381E-07 | NA |
| CWN-1::mNG | *control* | **mean intensity** | **15.006468** | **18.397206** | **20.516777** | **25.345054** | **24.754023** | **44.508472** |
| | | difference with next cell | -3.390738 | -2.119570 | -4.828277 | 0.591031 | -19.754449 | NA |
| | | paired t-test p-value | 0.000768 | 0.026574 | 0.008702 | 0.824677 | 1.7232E-08 | NA |
| | *cam-1* | **mean intensity** | **11.212148** | **13.090992** | **14.352518** | **16.982342** | **19.532900** | **30.200781** |
| | | difference with next cell | -1.878844 | -1.261527 | -2.629824 | -2.550558 | -10.667881 | NA |
| | | paired t-test p-value | 0.000436 | 0.007882 | 0.000182 | 0.005975 | 1.7467E-07 | NA |

**Table S3**

| **CAM-1::mNG intensity average per cell** | | | | | | |
|---|---|---|---|---|---|---|
| | V1 | V2 | V3 | V4 | V5 | V6 |
| ***control*** | **60.651658** | **50.100813** | **42.415773** | **36.839898** | **22.520003** | **28.746436** |
| ***cwn-1; egl-20*** | **89.967254** | **68.994956** | **57.231002** | **41.020058** | **30.053753** | **31.857299** |
| difference with control | -29.315596 | -18.894143 | -14.815228 | -4.180160 | -7.533750 | -3.110863 |
| t-test p-value | 3.3232E-06 | 3.188E-07 | 9.4745E-11 | 0.128796 | 0.001108 | 0.124423 |
| ***cwn-1; egl-20 cwn-2*** | **68.685568** | **59.273300** | **53.055434** | **42.324569** | **27.650412** | **31.873137** |
| difference with control | -8.033910 | -9.172487 | -10.639661 | -5.484670 | -5.130409 | -3.126701 |
| t-test p-value | 0.192515 | 0.022864 | 0.000648 | 0.067766 | 0.060502 | 0.264255 |
| **MOM-5::mNG intensity average per cell** | | | | | | |
| | V1 | V2 | V3 | V4 | V5 | V6 |
| ***control*** | **22.636050** | **27.752667** | **20.267128** | **21.560729** | **17.694786** | **13.871805** |
| ***cwn-1; egl-20*** | **29.929724** | **31.887427** | **24.659666** | **23.332197** | **18.929680** | **14.990488** |
| difference with control | -7.293674 | -4.134760 | -4.392537 | -1.771468 | -1.234894 | -1.118683 |
| t-test p-value | 2.2016E-06 | 0.000947 | 0.000131 | 0.183762 | 0.425404 | 0.133494 |
| ***control*** | **36.564263** | **42.982062** | **30.325845** | **31.676149** | **27.599700** | **17.900408** |
| ***cwn-1; egl-20 cwn-2*** | **31.188443** | **35.590080** | **27.461852** | **27.122233** | **23.302776** | **18.088667** |
| difference with control | 5.375820 | 7.391982 | 2.863993 | 4.553916 | 4.296924 | -0.188259 |
| t-test p-value | 0.002330 | 0.009347 | 0.000874 | 0.003873 | 0.080033 | 0.012551 |

**Table S4**

| mNG::MIG-5 intensity average per cell | | | | | | |
|---|---|---|---|---|---|---|
| | V1 | V2 | V3 | V4 | V5 | V6 |
| ***control*** | **16.265505** | **17.725742** | **17.818785** | **19.887060** | **22.393263** | **20.059526** |
| ***cwn-1; egl-20 cwn-2*** | **12.773407** | **14.701181** | **15.426054** | **18.048591** | **22.255418** | **17.732185** |
| difference with control | 3.492098 | 3.024561 | 2.392731 | 1.838469 | 0.137845 | 2.327342 |
| t-test p-value | 9.18059 E-08 | 0.000368 | 0.004898 | 0.095838 | 0.923739 | 0.010224 |
| **EGL-20::mNG intensity average per cell** | | | | | | |
| | V1 | V2 | V3 | V4 | V5 | V6 |
| ***control*** | **14.512751** | **20.379104** | **24.212777** | **33.215453** | **43.732358** | **48.812658** |
| ***dsh-1 dsh-2*** | **13.569336** | **18.069216** | **21.602779** | **28.410757** | **36.509846** | **34.082803** |
| difference with control | 0.943415 | 2.309888 | 2.609998 | 4.804696 | 7.222512 | 14.729855 |
| t-test p-value | 0.276571 | 0.073867 | 0.096467 | 0.010239 | 0.045490 | 1.75793 E-06 |
| **CWN-1::mNG intensity average per cell** | | | | | | |
| | V1 | V2 | V3 | V4 | V5 | V6 |
| ***control*** | **15.006468** | **18.397206** | **20.516777** | **25.345054** | **24.754023** | **44.508472** |
| ***dsh-1 dsh-2*** | **13.182266** | **15.038676** | **16.497323** | **17.302361** | **19.841744** | **22.835821** |
| difference with control | 1.824202 | 3.358531 | 4.019454 | 8.042693 | 4.912279 | 21.672651 |
| t-test p-value | 0.154702 | 0.022175 | 0.015174 | 0.000597 | 0.034424 | 1.22222 E-07 |
