## Supplemental files for "Reciprocal regulation of Wnt, receptor and Dvl localizations during cell polarization"

| Strain | Genotype |
|---|---|
| **CWN-1::mNG strains** | |
| HS3271 | ***cwn-1(cp205)*** *II [cwn-1::mNG^3XFlag.]; osIs33 IV [SCMp::mCherry::CAAX.]* |
| HS3340 | ***cwn-1(cp205)*** *II cam-1(gm122) II ; osIs33 IV* |
| HS3346 | *mom-5(ne12) I / hT2 I;III [qIs48];* ***cwn-1(cp205)*** *II ; osIs33 IV* |
| HS3404 | *lin-17(n3091) I ;* ***cwn-1(cp205)*** *II ; osIs33 IV* |
| HS3401 | *mom-5(os171) I [mom-5::AID::Clover];* ***cwn-1(cp205)*** *II cam-1(gm122) II ; osIs33 IV ; osIs114 [eft-3p::TIR-1 + coelRFP]* |
| HS3343 | ***cwn-1(cp205)*** *II ; egl-20(n585) IV osIs33 IV* |
| HS3571 | **cwn-1(cp205)** II dsh-2(or302) II dsh-1(ok1445) II / mIn1 II [mIs14 II dpy-10(e128) II ] ; osIs33 IV |
| **EGL-20::mNG strains** | |
| HS3292 | ***egl-20(cp221)*** *IV [egl-20::mNG^3XFlag] osIs33 IV* |
| HS3300 | *cam-1(gm122) II ;* ***egl-20(cp221)*** *IV osIs33 IV* |
| HS3299 | *mom-5(ne12) I / hT2 I;III [qIs48];* ***egl-20(cp221)*** *IV osIs33 IV* |
| HS3294 | *lin-17(n3091) I ;* ***egl-20(cp221)*** *IV osIs33 IV* |
| HS3399 | *mom-5(os171) I ; cam-1(gm122) II ;* ***egl-20(cp221)*** *IV osIs33 IV ; osIs118 X [eft-3p::TIR-1 + coelRFP]* |
| HS3907 | **egl-20(cp221)** IV osIs33 IV ; vang-1(tm1422) X |
| HS3342 | *cwn-1(ok546) II ;* ***egl-20(cp221)*** *IV osIs33 IV* |
| HS3576 | *dsh-2(or302) II dsh-1(ok1445) II / mIn1 II [dpy-10(e128) II mIs14 II ] ;* ***egl-20(cp221)*** *IV osIs33 IV* |
| **mNG::CWN-2 strains** | |
| HS3744 | ***cwn-2(os187)*** *IV [mNG::cwn-2] osIs33 IV* |
| HS3981 | *cam-1(gm122) II ;* ***cwn-2(os187)*** *IV osIs33 IV* |
| HS3985 | *mom-5(ne12) I / hT2 I;III ;* ***cwn-2(os187)*** *IV osIs33 IV* |
| HS4128 | *lin-17(n3091) I ;* ***cwn-2(os187)*** *IV osIs33 IV* |
| HS4424 | *lin-17(n3091) I mom-5(os171) I ;* ***cwn-2(os187)*** *IV osIs33 IV ; osIs253 V [Peft-3::TIR1(F79G)::mRuby::tbb-2 3'UTR]* |
| HS3983 | *cwn-1(ok546) II ; egl-20(os214) IV* ***cwn-2(os187)*** *IV osIs33 IV* |
| **CAM-1::mNG strains** | |
| HS3301 | ***cam-1(cp243)*** *II [cam-1::mNG-C1^3xFlag]); osIs33 IV* |
| HS3313 | ***cam-1(cp243)*** *II ; egl-20(n585) IV osIs33 IV* |
| HS3394 | *cwn-1(ok546) II* ***cam-1(cp243)*** *II ; osIs33 IV* |
| HS4466 | ***cam-1(cp243)*** *II ; cwn-2(ok895) IV osIs33 IV* |
| HS3400 | *cwn-1(ok546) II* ***cam-1(cp243)*** *II ; egl-20(n585) IV osIs33 IV* |
| HS4477 | *cwn-1(ok546) II* ***cam-1(cp243)*** *II ; egl-20(n585) IV cwn-2(ok895) IV osIs33 IV* |
| **MOM-5::mNG strains** | |
| HS3293 | ***mom-5(cp367)*** *I [mom-5::mNG-C1^3xFlag]; osIs33 IV* |

| | |
|---|---|
| HS3317 | ***mom-5(cp367)*** *I ; egl-20(n585) IV osIs33 IV* |
| HS4470 | ***mom-5(cp367)*** *I ; cwn-2(ok895) IV osIs33 IV* |
| HS3397 | ***mom-5(cp367)*** *I ; cwn-1(ok546) II ; egl-20(n585) IV osIs33 IV* |
| HS3315 | ***mom-5(cp367)*** *I ; cwn-1(ok546) II ; egl-20(n585) IV cwn-2(ok895) IV osIs33 IV* |
| **LIN-17::mNG strains** | |
| HS4488 | ***lin-17(os229)*** *I [lin-17::mNG::3xFlag ]; osIs33 IV* |
| **mNG::MIG-5 strains** | |
| HS4597 | ***mig-5 (cp385)*** *[mNG-GLO^AID::mig-5]; osIs33* |
| HS4602 | ***mig-5 (cp385);*** *egl-20 (n585) osIs33* |
| HS4617 | *cwn-1(ok546) II* ***mig-5 (cp385) II;*** *egl-20(n585) IV cwn-2(ok895) IV osIs33 IV* |
| **Colocalization strains** | |
| HS4613 | ***cam-1(os182)*** *[cam-1::mAID::mKate2] II;* ***egl-20(cp221)*** *IV; osIs276 [SCMp-BFP-CAAX]* |
| HS4614 | ***cam-1(os182)*** *II;* ***cwn-2(os187)*** *IV; osIs276* |
| HS4479 | ***mom-5(cp367)*** *I ;* ***cam-1(os182)*** *II ; osIs276* |
| HS4512 | ***mom-5(cp367)*** *I ; cwn-1(ok546) II* ***cam-1(os182)*** *II ; egl-20(n585) IV cwn-2(ok895) IV / tmC5 IV [tmIs1220 IV] ; osIs276* |
| HS4478 | ***cam-1(os182)*** *II* ***mig-5(os234)*** *[mNG::MIG-5] II ; osIs276* |
| HS4507 | *cwn-1(ok546) II* ***cam-1(os182)*** *II* ***mig-5(os234)*** *II ; egl-20(n585) IV cwn-2(ok895) IV / tmC5 IV [tmIs1220 IV ] ; osIs276* |
| **Other strains** | |
| HS4227 | *vang-1(tm1422) X ; wgIs559 [elt-3::TY1::EGFP::3xFLAG + unc-119(+)]* |

| Gene | Allele | Decription | Reference | doi |
|---|---|---|---|---|
| *cam-1* | *cp243* | *cam-1::mNG-C1^3xFlag* knock-in (C-terminal tag) | Heppert et al. 2018 | 10.1534/genetics.117.300487 |
| *cam-1* | *gm122* | Nonsense; lf (null) | Forrester et al. 1999 | 10.1038/23722 |
| *cam-1* | *os182* | *cam-1::mAID::mKate2* knock-in (auxin-inducible degron) | This study | |
| *cwn-1* | *cp205* | *cwn-1::mNG^3xFlag* knock-in (C-terminal tag) | same as *cp209* from Pani et al. 2023 | 10.1101/2023.05.03.539245 |
| *cwn-1* | *ok546* | Deletion; lf (null) | C. elegans Deletion Mutant Consortium 2012 | 10.1534/g3.112.003830 |
| *cwn-2* | *ok895* | Deletion; lf (null) | C. elegans Deletion Mutant Consortium 2012 | 10.1534/g3.112.003830 |
| *cwn-2* | *os187* | *mNG::cwn-2* knock-in (mNG inserted after signal sequence); functional | Sawa et al. 2026 | 10.1093/genetics/iyag004 |
| *dsh-1* | *ok1445* | Deletion; lf | C. elegans Deletion Mutant Consortium 2012 | 10.1534/g3.112.003830 |
| *dsh-2* | *or302* | Deletion; lf (maternal-effect embryonic lethal) | Hawkins et al. 2005 | 10.1016/j.ydbio.2005.05.024 |
| *egl-20* | *cp221* | *egl-20::mNG::3xFlag* knock-in (C-terminal tag); functional | Pani and Goldstein 2018 | 10.7554/eLife.38325 |
| *egl-20* | *n585* | Missense (conserved Cys); strong lf, behaves like null | Maloof et al. 1999 | 10.1242/dev.126.1.37 |
| *egl-20* | *os214* | Nonsense; lf (null) | This study | |
| *lin-17* | *n3091* | Nonsense; lf (null) | Sawa et al. 1996 | 10.1101/gad.10.17.2189 |
| *lin-17* | *os229* | *lin-17::mNG::3xFlag* knock-in (C-terminal tag) | This study | |
| *mig-5* | *cp385* | *mNG-GLO^AID*::mig-5* knock-in (N-terminal tag + auxin-inducible degron) | Heppert et al. 2018 | 10.1534/genetics.117.300487 |
| *mig-5* | *os234* | *mNG::mig-5* knock-in (N-terminal tag) | This study | |
| *mom-5* | *cp367* | *mom-5::mNG-C1^3xFlag* knock-in (C-terminal tag) | Heppert et al. 2018 | 10.1534/genetics.117.300487 |
| *mom-5* | *ne12* | lf (null); maternal-effect embryonic lethal | Rocheleau et al. 1997 | 10.1016/S0092-8674(00)80531-0 |
| *mom-5* | *os171* | *mom-5::mAID::mClover* knock-in (C-terminal, auxin-inducible degron) | Sawa et al. 2026 | 10.1093/genetics/iyag004 |
| *vang-1* | *tm1422* | Deletion; lf (null) | C. elegans Deletion Mutant Consortium 2012 | 10.1534/g3.112.003830 |
| | *osIs33* | IV; *SCMp::mCherry::CAAX* (seam cell membrane marker) (multicopy) | This study | |
| | *osIs114* | *eft-3p::TIR-1 + coel::RFP* (multicopy) | This study | |
| | *osIs118* | X; *eft-3p::TIR-1 + coel::RFP* (multicopy) | This study | |
| | osIs182 | V; *Peft-3::TIR1(F79G)::tbb-2 3'UTR* (AID2 system) (single copy) | This study | |
| | *osIs253* | V; *Peft-3::TIR1(F79G)::mRuby::tbb-2 3'UTR* (AID2 system) (single copy) | This study | |
| | *osIs276* | *SCMp::BFP::CAAX* (seam cell membrane marker) (multicopy) | This study | |
| | *wgIs559* | *elt-3::TY1::EGFP::3xFLAG + unc-119(+)* | Sarov et al. 2006; Zhong et al. 2010 | 10:839-44; 6(2):e1000848. |

| **EGL-20::mNG** | | | | | | | | | | | | | |
|---|---|---|---|---|---|---|---|---|---|---|---|---|---|
| | | V1 | | V2 | | V3 | | V4 | | V5 | | V6 | |
| Genotype | | A | P | A | P | A | P | A | P | A | P | A | P |
| *control 1* | mean | 12.7048182 | 16.3206842 | 17.2264966 | 23.5317112 | 20.6038625 | 27.8216919 | 29.797009 | 36.6338971 | 38.8849951 | 48.5797211 | 40.9764145 | 56.6489021 |
| | sd | 2.20111313 | 4.65187675 | 4.89474801 | 6.31261511 | 6.89256344 | 8.06631262 | 8.52258789 | 8.91429154 | 13.2494867 | 22.8617371 | 11.7246526 | 15.2659595 |
| | n | 28 | | 28 | | 28 | | 28 | | 28 | | 28 | |
| | paired t-test p value | 4.73521E-07 | | 6.51427E-11 | | 9.95288E-06 | | 0.000190707 | | 0.008132914 | | 1.54873E-06 | |
| *cam-1* | mean | 12.6035337 | 12.2608397 | 13.6190074 | 14.3036241 | 14.9694919 | 17.6157562 | 19.7062167 | 23.4177626 | 27.8401066 | 37.4009738 | 25.729108 | 40.8836261 |
| *compare with* | sd | 2.040576 | 1.90418791 | 1.69739096 | 1.9130294 | 1.8897332 | 3.91545974 | 3.98731323 | 4.37224668 | 7.51415529 | 12.2823899 | 6.02712247 | 12.6591227 |
| *control 1* | n | 27 | | 27 | | 27 | | 27 | | 27 | | 27 | |
| | paired t-test p value | 0.305170221 | | 0.081826457 | | 0.000170428 | | 6.46493E-05 | | 0.000198379 | | 5.91288E-08 | |
| *mom-5* | mean | 13.317249 | 17.7060701 | 16.3510735 | 20.6075826 | 19.7650013 | 25.0267925 | 31.4267953 | 33.9098213 | 37.357996 | 41.3960028 | 38.5369002 | 54.7539035 |
| *compare with* | sd | 3.36537756 | 7.02026561 | 4.03319229 | 5.80381219 | 5.12549999 | 6.91914623 | 7.63800428 | 9.61138618 | 10.8579014 | 17.3061807 | 10.3300593 | 17.2883788 |
| *control 1* | n | 34 | | 34 | | 33 | | 34 | | 34 | | 34 | |
| | paired t-test p value | 3.1606E-05 | | 5.10695E-07 | | 0.000181336 | | 0.182014346 | | 0.138443522 | | 5.42979E-07 | |
| *lin-17* | mean | 15.4067608 | 20.8878308 | 20.2021962 | 31.0580419 | 25.0443642 | 37.8659266 | 35.456117 | 49.6075831 | 44.0441581 | 55.9198166 | 42.4679895 | 54.689852 |
| *compare with* | sd | 3.97698999 | 6.00871313 | 5.69823812 | 9.63471833 | 6.36660148 | 10.9600305 | 10.6041334 | 14.4743506 | 12.9955125 | 11.2972962 | 10.1910435 | 13.2118558 |
| *control 1* | n | 32 | | 32 | | 32 | | 32 | | 32 | | 32 | |
| | paired t-test p value | 3.72843E-10 | | 1.09494E-09 | | 2.54727E-07 | | 5.89588E-06 | | 0.000286481 | | 4.23617E-08 | |
| *mom-5 (AID); cam-1* | mean | 11.9584121 | 12.2613328 | 13.1923979 | 13.822222 | 15.4360488 | 15.3189037 | 20.2404128 | 18.3595401 | 23.6142281 | 25.3866281 | 25.8997785 | 42.9339371 |
| *compare with* | sd | 1.43077278 | 1.29327623 | 1.91274134 | 1.91726274 | 1.74700555 | 1.95351837 | 4.81697194 | 3.65336857 | 5.49146042 | 7.97578453 | 4.67067846 | 8.0258292 |
| *control 1* | n | 26 | | 26 | | 26 | | 26 | | 26 | | 26 | |
| | paired t-test p value | 0.280279502 | | 0.174449677 | | 0.833562755 | | 0.13433078 | | 0.288693136 | | 1.34037E-11 | |
| *cwn-1* | mean | 12.7681017 | 15.5486432 | 15.2779103 | 22.9177425 | 19.2791742 | 27.616111 | 26.0887371 | 35.216864 | 31.5374401 | 40.0105655 | 37.9723977 | 53.4026811 |
| *compare with* | sd | 3.70654157 | 6.19968784 | 2.52980368 | 4.76851158 | 3.51133815 | 6.32018144 | 4.53473984 | 7.58121859 | 9.98904423 | 13.7092199 | 9.46501186 | 10.3718803 |
| *control 1* | n | 28 | | 28 | | 28 | | 28 | | 28 | | 28 | |
| | paired t-test p value | 9.92101E-05 | | 2.09531E-11 | | 6.43166E-07 | | 2.76948E-06 | | 0.001751024 | | 2.07178E-09 | |
| *dsh-1 dsh-2* | mean | 12.0950728 | 15.0435997 | 14.9376749 | 21.2007566 | 20.2898369 | 22.9157211 | 27.5425899 | 29.2789247 | 34.6234855 | 38.3962072 | 33.4212574 | 34.7443495 |
| *compare with* | sd | 3.19783936 | 4.42385395 | 3.73096299 | 5.85221944 | 6.04444831 | 5.8575055 | 8.48248979 | 6.8291131 | 12.392769 | 10.9735174 | 8.88898828 | 12.0725019 |
| *control 1* | n | 34 | | 35 | | 35 | | 35 | | 35 | | 34 | |
| | paired t-test p value | 4.25364E-05 | | 4.57268E-10 | | 0.006232555 | | 0.241036319 | | 0.098709107 | | 0.484620324 | |
| *control 2* | mean | 6.23900363 | 7.96248158 | 6.94839134 | 9.79434751 | 8.57860601 | 11.9473534 | 10.9862408 | 15.131745 | 14.6717778 | 18.2946974 | 16.7283955 | 26.8655149 |
| | sd | 1.13558547 | 1.71177033 | 1.11724788 | 2.02797096 | 2.16944725 | 3.07977843 | 3.79292528 | 4.07878082 | 4.96522954 | 5.9523102 | 3.92385985 | 6.9326083 |
| | n | 26 | | 26 | | 26 | | 26 | | 26 | | 26 | |
| | paired t-test p value | 1.79758E-05 | | 3.25438E-08 | | 5.9747E-05 | | 0.00014753 | | 0.000419681 | | 1.86644E-08 | |
| *vang-1* | mean | 7.77034381 | 8.62565014 | 8.7486395 | 12.0247394 | 10.4214351 | 14.0883078 | 12.2233416 | 17.0327501 | 17.2334858 | 21.8740768 | 19.2106877 | 27.412016 |
| *compare with* | sd | 1.72542539 | 1.60603201 | 2.53321566 | 3.18164188 | 2.08677448 | 3.63773504 | 4.85380488 | 5.92833344 | 6.5266252 | 8.29096369 | 4.38615729 | 6.40787056 |
| *control 2* | n | 28 | | 28 | | 28 | | 28 | | 28 | | 28 | |
| | paired t-test p value | 0.005234754 | | 5.15225E-07 | | 4.19779E-06 | | 9.04853E-05 | | 0.001209671 | | 3.30957E-08 | |

| **CWN-1::mNG** | | | | | | | | | | | | | |
|---|---|---|---|---|---|---|---|---|---|---|---|---|---|
| | | V1 | | V2 | | V3 | | V4 | | V5 | | V6 | |
| Genotype | | A | P | A | P | A | P | A | P | A | P | A | P |
| *control 1* | mean | 13.5467353 | 16.4662011 | 16.2258665 | 20.5685461 | 18.1399496 | 22.8936038 | 22.993651 | 27.6964566 | 26.0481726 | 23.4598732 | 34.4602746 | 54.556669 |
| | sd | 4.72845387 | 5.29102419 | 5.05421288 | 6.77623834 | 6.24343431 | 7.28362021 | 8.90127662 | 13.3456565 | 7.73011263 | 7.86215316 | 14.136294 | 20.7243732 |
| | n | 30 | | 30 | | 30 | | 30 | | 29 | | 30 | |
| | paired t-test p value | 2.05473E-05 | | 9.67268E-05 | | 0.000148115 | | 0.00136664 | | 0.029944733 | | 2.45074E-07 | |
| *control 2* | mean | 8.59995141 | 10.6237644 | 11.3074051 | 13.6391436 | 10.5859959 | 13.6487779 | 13.0440513 | 14.62265 | 17.597226 | 13.8092321 | 18.3124905 | 24.4024909 |
| | sd | 1.13772474 | 1.66629088 | 2.20301153 | 2.84905608 | 2.29710825 | 3.54238193 | 2.29579627 | 2.86189039 | 3.77255584 | 2.44268438 | 5.27983729 | 8.10736927 |
| | n | 35 | | 35 | | 35 | | 35 | | 35 | | 35 | |
| | paired t-test p value | 1.38378E-07 | | 9.76254E-05 | | 4.02641E-05 | | 0.005447346 | | 8.05621E-06 | | 0.000113863 | |
| *cam-1* | mean | 11.0544672 | 11.3698286 | 13.4844465 | 12.6975366 | 13.7062361 | 14.9988006 | 17.3126408 | 16.6520436 | 19.6557541 | 19.4100463 | 24.2543786 | 36.1471834 |
| *compare with* | sd | 1.78144353 | 1.90199219 | 3.09697571 | 2.31971181 | 2.7839209 | 4.18365446 | 5.78824946 | 4.31085004 | 6.89853438 | 5.8147222 | 8.34431666 | 15.0282479 |
| *control 1* | n | 28 | | 29 | | 29 | | 29 | | 29 | | 29 | |
| | paired t-test p value | 0.35311431 | | 0.109338701 | | 0.060974361 | | 0.439656887 | | 0.840954607 | | 1.27451E-05 | |
| *mom-5* | mean | 12.1875253 | 15.2552877 | 14.155785 | 16.2098828 | 15.9820127 | 18.6406389 | 21.1414194 | 22.4530195 | 23.2549237 | 21.7085555 | 27.6495559 | 38.9601051 |
| *compare with* | sd | 2.60394607 | 4.6886189 | 3.75127947 | 3.37519393 | 3.65140366 | 3.59297056 | 7.95696905 | 8.34560228 | 7.70577749 | 6.50959798 | 8.83294816 | 13.9990866 |
| *control 1* | n | 30 | | 30 | | 30 | | 30 | | 30 | | 30 | |
| | paired t-test p value | 0.000410534 | | 0.000841558 | | 0.000775152 | | 0.321910337 | | 0.110200456 | | 1.64444E-05 | |
| *lin-17* | mean | 8.33784281 | 10.9836224 | 11.5126223 | 15.5345642 | 11.7990703 | 15.6474975 | 14.330463 | 14.9160553 | 15.9859419 | 13.8332859 | 16.8536111 | 28.2599824 |
| *compare with* | sd | 8.36032592 | 11.0021794 | 11.5538033 | 15.5170123 | 11.8451872 | 15.7215285 | 14.3365244 | 14.8653189 | 16.0501225 | 13.9157863 | 16.8029042 | 28.0727686 |
| *control 2* | n | 39 | | 39 | | 39 | | 39 | | 39 | | 39 | |
| | paired t-test p value | 1.43213E-07 | | 1.15807E-07 | | 1.14514E-06 | | 0.424295267 | | 0.003162518 | | 1.77679E-10 | |
| *mom-5 (AID); cam-1* | mean | 11.5254418 | 12.5133676 | 14.3329555 | 13.938006 | 15.2903645 | 15.7874994 | 17.3005603 | 15.7616703 | 16.8827957 | 15.4800538 | 19.1489443 | 28.2063886 |
| *compare with* | sd | 2.38832236 | 3.17762614 | 3.65730457 | 3.84220171 | 4.04147681 | 4.17911912 | 3.61856803 | 3.77703877 | 4.47697864 | 3.40186435 | 4.14444862 | 9.24545749 |
| *control 2* | n | 28 | | 30 | | 30 | | 30 | | 30 | | 28 | |
| | paired t-test p value | 0.105090772 | | 0.529105773 | | 0.488513546 | | 0.031869452 | | 0.070798586 | | 3.97501E-06 | |
| *egl-20* | mean | 12.7108134 | 15.171718 | 14.3013962 | 19.5547056 | 15.3835359 | 19.3753005 | 20.6199999 | 24.1376831 | 21.3016992 | 21.9316346 | 29.2102337 | 41.460374 |
| *compare with* | sd | 4.36208313 | 4.88447316 | 4.41871051 | 7.92574853 | 3.81327146 | 4.70997425 | 4.81599994 | 5.8494603 | 6.22088261 | 6.95367376 | 7.680334 | 13.1723981 |
| *control 1* | n | 30 | | 30 | | 30 | | 30 | | 30 | | 30 | |
| | paired t-test p value | 1.42085E-06 | | 0.001747648 | | 0.000263318 | | 0.002887003 | | 0.56362079 | | 2.54757E-05 | |
| *dsh-1 dsh-2* | mean | 12.6663153 | 13.6982174 | 13.8806257 | 16.1967254 | 15.8095136 | 17.1851315 | 17.2230875 | 17.3816338 | 19.4923628 | 20.1911257 | 21.6617546 | 24.0098879 |
| *compare with* | sd | 4.0809838 | 4.66148503 | 4.62797721 | 5.97938467 | 4.97360864 | 6.2719946 | 5.37245476 | 4.74722584 | 9.00584425 | 9.75913701 | 8.81573967 | 10.9554353 |
| *control 1* | n | 21 | | 23 | | 23 | | 23 | | 23 | | 23 | |
| | paired t-test p value | 0.071456545 | | 0.006179072 | | 0.053566411 | | 0.878466797 | | 0.657494982 | | 0.136767647 | |

**mNG::CWN-2**

| Genotype | | V1 | | V2 | | V3 | | V4 | | V5 | | V6 | |
|---|---|---|---|---|---|---|---|---|---|---|---|---|---|
| | | A | P | A | P | A | P | A | P | A | P | A | P |
| *control 1* | mean | 24.8828828 | 34.3378533 | 20.542564 | 26.7991169 | 15.9889933 | 18.1169114 | 16.1811441 | 16.794421 | 14.5922078 | 13.7557205 | 13.2067798 | 13.5418326 |
| | sd | 6.03403071 | 8.95532135 | 4.34774738 | 7.3641365 | 2.95381849 | 3.80622668 | 2.17288194 | 2.28179379 | 2.76025941 | 2.02665031 | 2.14797971 | 1.64515766 |
| | n | 32 | | 32 | | 32 | | 32 | | 32 | | 32 | |
| | paired t-test p value | 9.57688E-07 | | 1.06566E-05 | | 0.003930045 | | 0.093587904 | | 0.040423557 | | 0.350636134 | |
| *cam-1* | mean | 14.9709259 | 15.6026746 | 16.1365301 | 16.4205021 | 16.5906841 | 17.651703 | 16.9618965 | 17.2444235 | 15.6855537 | 15.2612825 | 14.4991575 | 15.1282112 |
| *compare with* | sd | 2.18375875 | 2.1261971 | 1.91538835 | 2.11909817 | 2.27359564 | 2.85752363 | 2.16608188 | 2.33660939 | 2.31311672 | 1.49384603 | 1.80836884 | 2.29194965 |
| *control 1* | n | 31 | | 31 | | 31 | | 31 | | 31 | | 31 | |
| | paired t-test p value | 0.195070311 | | 0.463793922 | | 0.009545791 | | 0.557634028 | | 0.272570236 | | 0.051452107 | |
| *mom-5* | mean | 22.9196791 | 35.2403248 | 25.0926966 | 29.0036769 | 24.0441409 | 22.5495498 | 19.7281334 | 19.6700934 | 17.4884167 | 15.6355791 | 15.4816821 | 15.6212366 |
| *compare with* | sd | 5.27457039 | 9.77709087 | 5.30485308 | 5.95373775 | 4.33056321 | 3.636471 | 3.01145089 | 2.61331374 | 1.64719402 | 1.70090037 | 1.94983491 | 2.11718215 |
| *control 1* | n | 32 | | 32 | | 32 | | 31 | | 32 | | 32 | |
| | paired t-test p value | 2.74266E-09 | | 0.000578973 | | 0.10265037 | | 0.892412272 | | 1.06945E-05 | | 0.69314256 | |
| *lin-17* | mean | 29.9759562 | 39.9901389 | 20.4873888 | 25.4877558 | 17.7974801 | 18.5262897 | 15.8363312 | 16.7992075 | 14.907294 | 13.8466339 | 13.4077549 | 14.2149256 |
| *compare with* | sd | 8.10256508 | 10.5133068 | 4.65577511 | 4.74219309 | 3.44140229 | 3.74903151 | 2.45330069 | 3.26374973 | 2.99531676 | 2.17514998 | 1.99842551 | 2.67968076 |
| *control 1* | n | 36 | | 36 | | 36 | | 36 | | 36 | | 36 | |
| | paired t-test p value | 8.69768E-06 | | 4.94427E-08 | | 0.263953892 | | 0.028361474 | | 0.028404163 | | 0.042763224 | |
| *mom-5 (AID) lin-17* | mean | 21.1672452 | 33.0866341 | 20.0258353 | 21.2439522 | 20.2449331 | 17.121568 | 16.4278389 | 16.3797453 | 15.7543078 | 13.8701412 | 12.575423 | 13.0121602 |
| *compare with* | sd | 4.81954022 | 11.0382193 | 5.84398901 | 5.43490876 | 5.28000083 | 2.45317978 | 3.62700372 | 3.53531321 | 3.51418953 | 1.69035083 | 1.53767151 | 2.17482001 |
| *control 1* | n | 36 | | 36 | | 36 | | 36 | | 36 | | 36 | |
| | paired t-test p value | 2.26392E-08 | | 0.196293859 | | 0.000403332 | | 0.929491122 | | 0.00090082 | | 0.172610839 | |
| *cwn-1; egl-20* | mean | 44.132736 | 47.0982425 | 27.5114995 | 35.5629045 | 21.4043885 | 23.8464765 | 23.2340093 | 24.9477524 | 17.5868455 | 16.1262895 | 14.3128726 | 15.411447 |
| *compare with* | sd | 30.9380359 | 12.4834167 | 6.3742318 | 7.79974013 | 5.23630013 | 6.50944307 | 3.73163287 | 4.45321122 | 2.61352721 | 2.2924321 | 2.44365809 | 2.57466763 |
| *control 1* | n | 35 | | 35 | | 35 | | 35 | | 35 | | 35 | |
| | paired t-test p value | 0.494738888 | | 1.04945E-07 | | 0.021349271 | | 0.080722911 | | 0.00653207 | | 0.005622894 | |

**CAM-1::mNG**

| Genotype | | V1 | | V2 | | V3 | | V4 | | V5 | | V6 | |
|---|---|---|---|---|---|---|---|---|---|---|---|---|---|
| | | A | P | A | P | A | P | A | P | A | P | A | P |
| *control 1* | mean | 81.5164383 | 92.1368357 | 60.3953051 | 77.0640914 | 49.615063 | 68.3258459 | 37.355034 | 59.1267348 | 23.0526543 | 32.9935945 | 30.7094769 | 49.5474433 |
| | sd | 31.3955139 | 30.5063222 | 14.1246068 | 24.4480533 | 13.8799906 | 16.0521143 | 10.2210867 | 15.369027 | 6.5436447 | 6.9927216 | 10.00994 | 18.2452256 |
| | n | 35 | | 35 | | 35 | | 25 | | 16 | | 35 | |
| | paired t-test p value | 0.000596762 | | 6.32965E-06 | | 5.49369E-08 | | 3.09643E-07 | | 3.11825E-05 | | 1.32252E-06 | |
| *control 2* | mean | 55.7313834 | 65.5719318 | 42.4724652 | 57.7291597 | 38.7173185 | 46.1142282 | 27.9988671 | 45.6809291 | 20.3950272 | 24.6449792 | 21.1288406 | 36.3640321 |
| | sd | 23.2113876 | 21.5747739 | 9.9762376 | 15.253896 | 7.77450033 | 9.68985836 | 7.15978352 | 15.0832475 | 6.06844977 | 6.87990458 | 6.950459 | 13.115138 |
| | n | 35 | | 36 | | 36 | | 22 | | 14 | | 36 | |
| | paired t-test p value | 4.64099E-05 | | 1.92726E-10 | | 0.000110724 | | 7.24546E-07 | | 0.042231107 | | 1.88837E-09 | |
| *egl-20* | mean | 94.6403147 | 107.848517 | 68.8033483 | 87.1343106 | 57.1708741 | 64.1455523 | 47.6637019 | 49.5161492 | 32.5217998 | 33.4044061 | 37.2011767 | 45.2705244 |
| *compare with* | sd | 29.5936713 | 36.0144627 | 14.3061493 | 24.6083131 | 10.2525902 | 18.0628812 | 10.6095176 | 14.9215907 | 13.7654676 | 12.021055 | 17.2200318 | 18.2543577 |
| *control 1* | n | 32 | | 32 | | 31 | | 25 | | 15 | | 31 | |
| | paired t-test p value | 0.004277933 | | 4.87188E-06 | | 0.018431443 | | 0.544072289 | | 0.796790968 | | 0.003510559 | |
| *cwn-1* | mean | 70.0337906 | 88.6051722 | 55.2773287 | 67.5079404 | 47.8575821 | 57.147153 | 37.8804569 | 42.070038 | 24.809332 | 35.6707645 | 30.3505041 | 46.8872658 |
| *compare with* | sd | 29.2926227 | 25.1759501 | 12.9818642 | 18.2931342 | 7.33968616 | 10.172147 | 9.48628423 | 9.88578811 | 4.75418346 | 6.85832953 | 7.9579006 | 16.7162044 |
| *control 2* | n | 32 | | 33 | | 33 | | 22 | | 11 | | 32 | |
| | paired t-test p value | 0.000161483 | | 6.39302E-06 | | 3.13487E-05 | | 0.120860393 | | 0.000573913 | | 1.40062E-06 | |
| *cwn-2* | mean | 43.3548467 | 65.55901 | 42.3744286 | 64.6997822 | 40.7184122 | 56.3264849 | 31.2543625 | 47.2487832 | 26.1620307 | 31.2282223 | 24.4348732 | 30.5636709 |
| *compare with* | sd | 16.4083186 | 20.8926654 | 10.2994102 | 16.0488725 | 7.53967484 | 15.443101 | 8.67466741 | 14.6222001 | 7.43228001 | 11.0431703 | 11.7523559 | 9.19841388 |
| *control 2* | n | 29 | | 31 | | 31 | | 21 | | 10 | | 30 | |
| | paired t-test p value | 1.53153E-08 | | 5.1503E-12 | | 7.95001E-08 | | 0.000144848 | | 0.26781111 | | 0.006141054 | |
| *cwn-1; egl-20* | mean | 80.5383338 | 99.3961737 | 60.7793669 | 77.2105449 | 57.6892647 | 56.7727387 | 41.8510481 | 40.1890674 | 33.4027838 | 26.7047216 | 30.4250911 | 33.2895065 |
| *compare with* | sd | 32.4586478 | 28.39634 | 15.9630428 | 19.713369 | 11.3957276 | 10.724263 | 8.82767217 | 8.50944835 | 8.69722622 | 4.53624093 | 9.11972306 | 9.06781578 |
| *control 2* | n | 36 | | 36 | | 36 | | 23 | | 15 | | 31 | |
| | paired t-test p value | 0.000365514 | | 6.20079E-07 | | 0.665594077 | | 0.341479919 | | 0.005848663 | | 0.127203953 | |
| *cwn-1; egl-20 cwn-2* | mean | 64.9322923 | 72.4388435 | 55.2499433 | 63.296656 | 51.1235578 | 54.9873102 | 43.4263844 | 41.2227527 | 27.71858 | 27.5822444 | 31.893241 | 31.8530339 |
| *compare with* | sd | 29.3454389 | 25.459626 | 23.4266338 | 17.6870393 | 14.9831725 | 15.533801 | 11.1221438 | 11.1267943 | 8.06894984 | 11.5135572 | 13.3184717 | 13.239665 |
| *control 2* | n | 25 | | 31 | | 31 | | 26 | | 15 | | 25 | |
| | paired t-test p value | 0.153409742 | | 0.01685105 | | 0.018308005 | | 0.255381775 | | 0.96141419 | | 0.98735921 | |

| **MOM-5::mNG** | | | | | | | | | | | | | |
|---|---|---|---|---|---|---|---|---|---|---|---|---|---|
| | | V1 | | V2 | | V3 | | V4 | | V5 | | V6 | |
| Genotype | | A | P | A | P | A | P | A | P | A | P | A | P |
| *control 1* | mean | 31.8458392 | 41.2826873 | 36.657432 | 49.306693 | 25.9416774 | 34.7100122 | 29.3939715 | 33.9583267 | 19.9112989 | 35.2881011 | 15.9766948 | 19.8241211 |
| | sd | 10.7928784 | 14.4374728 | 9.49035045 | 15.0803224 | 6.43404021 | 10.2458951 | 9.27980866 | 12.1379442 | 5.06807248 | 9.8678317 | 4.40196135 | 6.57025669 |
| | n | 26 | | 26 | | 26 | | 14 | | 13 | | 26 | |
| | paired t-test p value | 0.000626903 | | 5.97008E-07 | | 1.62043E-05 | | 0.187138689 | | 1.30213E-05 | | 0.001907371 | |
| *control 2* | mean | 20.5696776 | 24.7024216 | 23.522766 | 31.982568 | 17.0814133 | 23.4528432 | 18.7150442 | 23.8071273 | 15.5611559 | 19.8284155 | 11.1933617 | 16.5502483 |
| | sd | 4.69149965 | 6.72110017 | 3.76767066 | 6.54417279 | 3.76866364 | 6.79754909 | 3.58698246 | 5.96085575 | 4.53582485 | 5.24966272 | 2.16114023 | 4.88503144 |
| | n | 28 | | 30 | | 30 | | 19 | | 12 | | 29 | |
| | paired t-test p value | 4.91137E-06 | | 3.37248E-07 | | 7.61064E-07 | | 0.001751082 | | 0.022513646 | | 7.96093E-07 | |
| *egl-20* | mean | 36.3570849 | 54.5429556 | 39.5044654 | 49.4320376 | 31.4700518 | 36.4197507 | 34.5055529 | 37.6099019 | 21.5648448 | 21.7727052 | 20.8872788 | 22.4718427 |
| *compare with* | sd | 9.7572721 | 11.519486 | 10.012862 | 10.7908135 | 5.70054412 | 7.08213481 | 9.5774255 | 9.01037525 | 4.29617733 | 2.66791902 | 4.26518203 | 5.84316509 |
| *control 1* | n | 27 | | 27 | | 27 | | 17 | | 10 | | 23 | |
| | paired t-test p value | 5.17312E-08 | | 8.90452E-05 | | 0.001035132 | | 0.245799098 | | 0.883976387 | | 0.107021171 | |
| *cwn-2* | mean | 21.3174122 | 31.3436246 | 26.5131784 | 37.1795436 | 20.6237622 | 28.7098697 | 23.5196213 | 27.177536 | 18.014293 | 25.2929624 | 12.5822828 | 12.8655405 |
| *compare with* | sd | 6.71553699 | 10.6992742 | 7.50746003 | 10.237887 | 4.44410606 | 8.20580223 | 8.0551984 | 10.2173242 | 3.91233053 | 5.55172455 | 3.39960129 | 4.06794141 |
| *control 2* | n | 37 | | 38 | | 38 | | 21 | | 18 | | 37 | |
| | paired t-test p value | 6.32582E-10 | | 1.31262E-08 | | 1.45054E-07 | | 0.142240457 | | 4.43197E-05 | | 0.687700224 | |
| *cwn-1; egl-20* | mean | 27.6486737 | 32.2107741 | 30.262421 | 33.5124322 | 22.7719914 | 26.54734 | 25.2125629 | 21.4518314 | 20.1139748 | 17.745385 | 15.2145811 | 14.7663943 |
| *compare with* | sd | 6.41807182 | 7.13679158 | 6.21865985 | 7.3570975 | 4.51762559 | 4.84531856 | 5.85895595 | 4.73518808 | 4.60034179 | 2.74872451 | 3.20719287 | 3.16704018 |
| *control 2* | n | 29 | | 33 | | 33 | | 23 | | 15 | | 27 | |
| | paired t-test p value | 0.008534047 | | 0.03292472 | | 0.001128418 | | 0.002560273 | | 0.028360557 | | 0.566618218 | |
| *cwn-1; egl-20 cwn-2* | mean | 30.3632945 | 32.0135911 | 35.7017653 | 35.4783951 | 26.204158 | 28.7195451 | 30.5468536 | 23.6976125 | 24.080507 | 22.5250445 | 17.4516007 | 18.725733 |
| *compare with* | sd | 14.5014939 | 12.6654679 | 14.4219879 | 16.0004891 | 8.36156756 | 11.2732412 | 8.3309327 | 5.35618445 | 11.7584915 | 10.9984352 | 8.60257992 | 8.2280892 |
| *control 1* | n | 28 | | 28 | | 28 | | 16 | | 16 | | 28 | |
| | paired t-test p value | 0.36000599 | | 0.903355511 | | 0.025566013 | | 0.001703843 | | 0.239503138 | | 0.239103265 | |

| **LIN-17::mNG** | | | | | | | | | | | | | |
|---|---|---|---|---|---|---|---|---|---|---|---|---|---|
| | | V1 | | V2 | | V3 | | V4 | | V5 | | V6 | |
| Genotype | | A | P | A | P | A | P | A | P | A | P | A | P |
| *control* | mean | 15.9230408 | 19.6595515 | 25.6184301 | 26.8002804 | 36.0119395 | 45.4213854 | 46.6808351 | 41.6996249 | 83.1132294 | 78.6913308 | 55.9626418 | 87.4887887 |
| | sd | 3.49742378 | 4.70908599 | 4.07161557 | 4.69805326 | 6.48328694 | 6.95570063 | 10.1146228 | 9.5066281 | 14.8769253 | 14.125872 | 13.9180372 | 28.012354 |
| | n | 29 | | 29 | | 29 | | 15 | | 13 | | 29 | |
| | paired t-test p value | 4.75572E-06 | | 0.072561663 | | 1.2298E-10 | | 0.075554478 | | 0.207537557 | | 3.20057E-10 | |

| **mNG::MIG-5** | | | | | | | | | | | | | |
|---|---|---|---|---|---|---|---|---|---|---|---|---|---|
| | | V1 | | V2 | | V3 | | V4 | | V5 | | V6 | |
| Genotype | | A | P | A | P | A | P | A | P | A | P | A | P |
| *control 1* | mean | 14.8265179 | 17.7044925 | 16.1499065 | 19.301577 | 16.2687027 | 19.3688676 | 19.5628723 | 20.2112472 | 20.7503442 | 24.0361808 | 17.2984193 | 22.8206331 |
| | sd | 1.94628163 | 2.28450254 | 2.37596231 | 3.46381104 | 2.41751614 | 3.94631937 | 4.41576239 | 4.99470809 | 5.31997004 | 6.69921818 | 3.27406217 | 4.74482977 |
| | n | 29 | | 29 | | 29 | | 29 | | 29 | | 29 | |
| | paired t-test | 1.79136E-07 | | 1.17112E-06 | | 2.33437E-07 | | 0.356362391 | | 0.000878745 | | 4.14125E-08 | |
| *egl-20* | mean | 13.8767446 | 16.2704056 | 15.3355735 | 19.1090443 | 16.4677163 | 19.4314266 | 22.5412459 | 20.6723846 | 23.2522677 | 20.3561376 | 17.0890613 | 21.844943 |
| | sd | 1.68457132 | 2.80491673 | 2.17577342 | 3.77596762 | 2.57874661 | 3.19002636 | 7.09841649 | 3.68133816 | 4.75808958 | 4.16237611 | 3.21868612 | 4.94334724 |
| | n | 26 | | 26 | | 26 | | 26 | | 26 | | 26 | |
| | paired t-test | 4.30495E-05 | | 6.91485E-07 | | 4.39461E-05 | | 0.139822712 | | 0.00233049 | | 1.47942E-06 | |
| *cwn-1; egl-20 cwn-2* | mean | 12.3652952 | 13.1815182 | 14.5838268 | 14.8185352 | 14.9568048 | 15.895303 | 18.2108035 | 17.8863779 | 22.4736587 | 22.0371771 | 17.1863587 | 18.2780106 |
| | sd | 2.34821478 | 2.82095143 | 3.79952393 | 3.21896227 | 3.1216009 | 3.65710494 | 4.57293708 | 3.6855566 | 6.63467806 | 5.49765833 | 3.30282439 | 3.41709093 |
| | n | 29 | | 29 | | 29 | | 29 | | 29 | | 29 | |
| | paired t-test | 0.020835182 | | 0.536724149 | | 0.042515752 | | 0.526547345 | | 0.689398213 | | 0.033066846 | |

| CAM-1::mNG intensity average per cell | | Same as Table S3 | | | | |
|---|---|---|---|---|---|---|
| | V1 | V2 | V3 | V4 | V5 | V6 |
| control | 60.6516576 | 50.1008125 | 42.4157734 | 36.8398981 | 22.5200032 | 28.7464363 |
| *cwn-1; egl-20* | 89.9672538 | 68.9949559 | 57.2310017 | 41.0200578 | 30.0537527 | 31.8572988 |
| difference with control | -29.315596 | -18.894143 | -14.815228 | -4.1801597 | -7.5337495 | -3.1108625 |
| t-test | 3.3232E-06 | 3.188E-07 | 9.4745E-11 | 0.12879565 | 0.00110797 | 0.12442344 |
| *cwn-1; egl-20 cwn-2* | 68.6855679 | 59.2732997 | 53.055434 | 42.3245686 | 27.6504122 | 31.8731374 |
| difference with control | -8.0339103 | -9.1724872 | -10.639661 | -5.4846704 | -5.130409 | -3.1267011 |
| t-test | 0.1925151 | 0.02286432 | 0.00064751 | 0.06776614 | 0.06050153 | 0.26425456 |
| MOM-5::mNG intensity change per cell | | | | | | |
| | V1 | V2 | V3 | V4 | V5 | V6 |
| control | 22.6360496 | 27.752667 | 20.2671282 | 21.5607289 | 17.6947857 | 13.871805 |
| *cwn-1; egl-20* | 29.9297239 | 31.8874266 | 24.6596657 | 23.3321971 | 18.9296799 | 14.9904877 |
| difference with control | -7.2936743 | -4.1347596 | -4.3925375 | -1.7714682 | -1.2348942 | -1.1186828 |
| t-test | 2.2016E-06 | 0.00094735 | 0.00013062 | 0.18376198 | 0.42540407 | 0.13349433 |
| control | 36.5642632 | 42.9820625 | 30.3258448 | 31.6761491 | 27.5997 | 17.900408 |
| *cwn-1; egl-20 cwn-2* | 31.1884428 | 35.5900802 | 27.4618516 | 27.1222331 | 23.3027757 | 18.0886668 |
| difference with control | 5.37582042 | 7.39198227 | 2.86399322 | 4.55391606 | 4.29692426 | -0.1882589 |
| t-test | 0.00233001 | 0.00934688 | 0.00087432 | 0.0038733 | 0.08003324 | 0.01255075 |
